# A molecular glue in plants: The lectin domain of LecRK-I.9 creates persistent plasma membrane – cell wall connections

**DOI:** 10.1101/2025.08.14.670329

**Authors:** Denise S. Arico, Ugo Segura, Annalisa Bellandi, Claire Lionnet, Marjolaine Martin, Olivier Hamant, Hervé Canut

**Affiliations:** Laboratoire de Reproduction et Développement des Plantes, ENS de Lyon, UCBL, INRAE, CNRS, 46 Allée d’Italie, 69364 Lyon Cedex 07, France; Laboratoire de Recherche en Sciences Végétales, Université de Toulouse, CNRS, UPS, Toulouse INP, F-31320, Auzeville-Tolosane, France

**Author notes:** **AUTHOR CONTRIBUTIONS:** Conceptualization, D.S.A., O.H., and H.C.; methodology, D.S.A., A.B., and C.L.; Investigation, D.S.A., U.S.; writing—original draft, D.S.A. and O.H.; writing—review & editing, D.S.A., A.B., O.H., and H.C.; funding acquisition, O.H.; resources, D.S.A. and M.M.; supervision, H.C., O.H., and D.S.A. **DECLARATION OF INTERESTS:** The authors declare no competing interests.

**Keywords:** Cell wall – plasma membrane adhesion, Receptor-like kinase, Lectin domain, LecRK-I.9, plasma membrane contact sites, protein clusters, cell wall, osmotic stress

## Abstract

Water stress challenges plasma membrane – wall attachment. When plants are exposed to strong hyperosmotic stress, water exits the cell leading to plasmolysis, where the plasma membrane detaches partially from the cell wall and reveals discrete and persistent attachment sites in the form of Hechtian strands. Although these structures have been observed in several species for more than a century, the molecular bases behind plasma membrane – wall attachment remains elusive. Through a screen of candidate proteins, we reveal that the overexpression of two lectin receptor-like proteins (LecRK-I.9 and LecTM) increases the density of plasma membrane – wall connections in *N. Benthamiana*. The extracellular lectin domain of LecRK-I.9 was able to remain in the wall during plasmolysis. Conversely, deletion of the lectin domain in the LecRK-I.9 overexpressor restored the density of Hechtian strands to WT levels. In Arabidopsis, upon hyperosmotic stress, LecRK-I.9 formed largely immobile clusters, whereas lectin-deleted versions of LecRK-I.9 clusters were mobile. Cluster density correlated with predicted tensile stress levels and mechanical reinforcement in the wall before plasmolysis, consistent with a scenario in which attachment of the lectin domain also reflects the wall properties. Last, while overexpressing LecRK-I.9 conferred resistance to low water potential conditions, deletion of the lectin domain in the LecRK-I.9 overexpressor restored a WT response to water stress. Altogether, this demonstrates that the lectin domain of LecRK-I.9 creates persistent plasma membrane – wall attachment sites, with physiological relevance for plant resistance to water stress.

**SIGNIFICANCE STATEMENT:** As observed since the mid-19^th^ century, when placed in hyperosmotic conditions, plant cells undergo plasmolysis and form thin membraneous threads called Hechtian strands. These structures reflect the presence of persistent plasma membrane – cell wall attachment sites. Yet, the molecular components behind these sites are unknown. Through a candidate screen approach, we identify and demonstrate that the lectin domain of a receptor-like kinase provides such molecular glue. We also show that the clustering of the receptor is patterned and correlate with mechanically reinforced cell walls. Conversely, we find that the promotion of attachment increases resistance to water stress. This work revisits an old plant cell biology classic, and opens new avenues of research for plant adaptation to their environment.

## INTRODUCTION

Plants are continuously exposed to endogenous and exogenous mechanical stress, stemming from factors such as high turgor pressure (1, 2) and mechanical perturbations from the environment (3, 4). To resist such stress, plant cells reinforce their walls. Beyond mechanical stress, plants are also exposed to biochemical signals, which must also be transduced through the cell wall to the cytoplasm(5). Consistently, there is increasing evidence that the cell wall – plasma membrane nexus can function as a structural and a signaling hub for plant integrity and development (6, 7). For instance, cellulose synthesis involves a protein complex, which directionality is biased by stress-dependent orientation of cortical microtubules (8–10), and which activity is influenced by osmotic conditions and hormone signaling (11–14). Furthermore, the plasma membrane hosts a long list of receptor-like kinases, with extracellular domains that might anchor them into the cell wall.

Plasma membrane – cell wall connections are evidenced during plasmolysis. When plants are exposed to strong hyperosmotic stress (e.g. drought, high soil salinity, or freezing), water flows out of the cell leading to the retraction of the plasma membrane that detaches partially from the cell wall. This detachment reveals discrete and persistent sites, where the cell wall remains connected to the plasma membrane. The resulting membrane strands are called Hechtian strands (15). While many of the Hechtian strands are linked to plasmodesmata, many others are independent from plasmodesmata, thus revealing that plasma membrane – cell wall connections can, on their own, provide adhesive hotspots at the wall (Figure 1a)(15). The identity of the molecular linker in these attachment sites is unknown.

**Figure 1.**
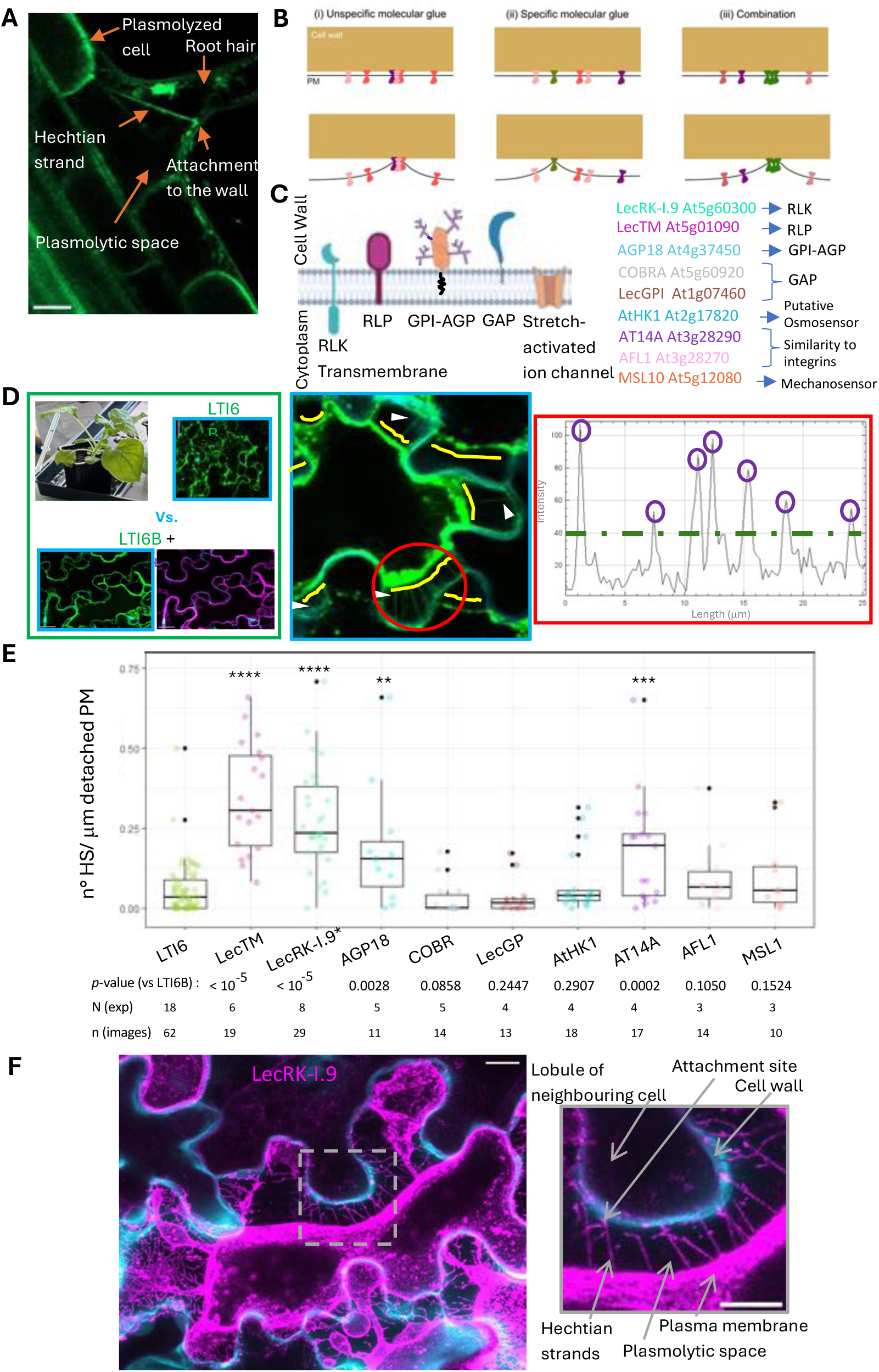
Screen for proteins promoting persistent plasma membrane – cell wall connections in *Nicotiana benthamiana*. **A**. Confocal image of Arabidopsis root expressing *p35S::eGFP-LTI6B*, that shows a plasmodesmata-unrelated Hechtian strand traversing the plasmolytic space in a root hair cell plasmolyzed with sorbitol 0.6M. Scale bar = 10μm. **B.** Schematic representation of the working hypothesis, illustrating potential scenarios where plasma membrane proteins act as molecular glue between the plasma membrane and the cell wall. **C.** Scheme (Illustrated with BioRender) and brief description of candidate proteins located at the plasma membrane – wall interface. RLK: Receptor Like Kinase. RLP: Receptor Like Protein. GPI-AGP: Glycosyl-Phosphatidyl-Inositol – ArabinoGalactan Protein. GAP: GPI- Anchored Protein. **D.** Transient transformation of *N. benthamiana* followed by plasmolysis of pavement cells with glycerol 10%, and quantification of density of Hechtian strands labeled with *LTI6B-mCitrine*. Proceedings for the screening detailed in Methods section. **E.** Boxplot showing the density of Hechtian strands (HS), expressed as number of HS per μm of detached plasma membrane (PM), for each tested candidate compared to the control LTI6B. Boxes indicate the interquartile range (IQR), the line inside each box represents the median, whiskers show the data range within 1.5×IQR, and black dots indicate outliers. Statistical significance was determined using the Kruskal–Wallis test followed by Dunn’s post hoc test with Benjamini–Hochberg correction. Sample size (N = number of experiments, n = number of images with 1 -3 cells averaged) and *p*-values are indicated below the plot. Asterisks denote statistically significant differences compared to the LTI6B control. **F.** Confocal image showing a *N. benthamiana* pavement cell (at the middle-plane) expressing LecRK-I.9*-TagRFP (magenta), after plasmolysis with glycerol 20%. The cell walls were stained with calcofluor (cyan). Scale bar = 12 μm. Inset showing the Hechtian strands in detail. Scale bar = 10μm.

Several scenarios may be envisioned (Figure 1b). Protein diffusion at the plasma membrane is often constrained by interactions with the cell wall or cytoskeleton, leading to the formation of nanodomains—local accumulation of molecules within a membrane plane at the nanoscale (16)—which have been implicated in processes such as signaling and mechanosensing (17, 18). Thus, one may hypothesize that the clustering of membrane proteins is sufficient to provide increased adhesion at discrete hotspots, with no protein specifically acting as a linker (e.g. cooperative behavior among proteins with low affinity binding for the cell wall or projecting into the apoplast). Second, certain classes of proteins could bind cell wall components with high affinity and provide strong adhesion at discrete hotspots. Third, a combination of membrane proteins with different binding affinities for cell wall components would be necessary to provide such persistent adhesion sites. In the end, this comes to the hypothesis that plasma membrane - wall adhesion is an emergent property arising from the spatial organization of membrane proteins and their wall adhesion strength.

In order to test whether such wall-adhesive membrane proteins exist or not, we initiated a screening of *Arabidopsis thaliana* proteins located at the plasma membrane- cell wall interface. We found that the lectin domain of receptor-like proteins has such a property.

## RESULTS

### The overexpression of LecRK-I.9* and LecTM increases the number of plasma membrane – cell wall connections in plasmolyzed pavement cells of *Nicotiana benthamiana*

To identify whether plasma membrane - cell wall connections are under the control of specific molecular factors, we selected a short list of *Arabidopsis thaliana* candidate proteins documented to have either the ability to interact with some components of the wall or to contribute to the transmission of mechanical signals (Figure 1c). We reasoned that mechanosensing factors involve structural components that can transmit physical signals from the wall to the membrane, and thus are ideal candidates to reveal persistent adhesion hotspots at the cell wall – plasma membrane nexus. We targeted candidates with transmembrane domains and extracellular domains that may interact with the wall(19, 20), such as the Legume-type (L-type) Lectin Receptor-like kinase LecRK-I.9 (At5g60300) and a receptor-like protein LecTM (At5g01090). In addition, we considered candidate proteins with reported partial sequence similarities to integrins that connect the extracellular matrix with the actin cytoskeleton in mammalian cells (21, 22), such as AT14A (At3g28290) and AFL1 (At3g28270). We also included extracellular proteins bound to the plasma membrane with a Glycosyl-phosphatidyl-inositol (GPI) anchor (23, 24), such as AGP18 (At4g37450), COBRA (At5g60920) and LecGPI (At1g07460). Finally, we chose the well-established mechanosensory (25) MSL10 (At5g12080) and a protein described as a putative osmosensor (26, 27) AtHK1 (At2g17820) (Figure 1c). Note that we tested a kinase-dead version of LecRK-I.9 (noted LecRK-I.9*, see Methods section and Abbreviations), as the overexpression of native LecRK-I.9 in Arabidopsis appeared to generate phenotypic defects which impaired the analysis of its localization (Figure S1). This is also consistent with the putative toxicity of LecRK-I.8-GFP overexpression, where very low level of LecRK-I.8-GFP in constitutively overexpressing lines was detected only in three out of eleven independent lines(28). The LecRK-I.9* kinase-dead version is a full-length version of LecRK-I.9 with 5 alanine substitutions at positions 350aa – 354aa within the ATP binding motif LGKGG.

We hypothesized that, upon plasmolysis, a cell overexpressing a protein that generates persistent attachment sites to the wall would exhibit an increased number of Hechtian strands compared to a Wild-Type (WT) cell. To do so, we developed a simple pipeline to quantify the number of Hechtian strands (labeled with the membrane marker LTI6B) per μm of detached plasma membrane (density of Hechtian strands), in plasmolyzed pavement cells from *N. benthamiana* overexpressing each candidate compared to WT cells (Figure 1d). Despite the high variability and low density of Hechtian strands, two proteins stood out: the overexpression of LecTM and of the kinase-dead version of LecRK-I.9 (LecRK-I.9*) strongly increased the density of Hechtian strands in plasmolyzed pavement cells of *N. benthamiana* (Figure 1e). Note that the overexpression of AGP18 and AT14A also increased the density of Hechtian strands but the effect was milder (Figure 1e).

LecRK-I.9 can interact *in vitro* with the *Phytophthora infestans* effector IPI-O (29), which is an RGD-containing peptide that was reported to disrupt plasma membrane – cell wall adhesions in Arabidopsis (30). This, together with our screen results highlighted LecRK-I.9 as the most attractive candidate to act as a plasma membrane – cell wall glue (Figure 1f).

### The overexpression of LecRK-I.9* specifically increases the plasma membrane – cell wall connections related to Hechtian strands and Hechtian reticulum

In order to test if the overexpression of LecRK-I.9* also increased the connection between the plasma membrane and the wall apart from Hechtian strands, we developed a pipeline to quantify the percentage of plasma membrane that remained attached to the wall after plasmolysis in cells overexpressing LecRK-I.9* vs. LTI6B. First, we compared the proportion of the plasma membrane length that remained attached to the wall relative to the total cell perimeter at the cell middle plane, in cells exhibiting the same degree of plasmolysis. A plasmolysis index was calculated as the difference in cell area before and after plasmolysis, normalized to the pre-plasmolysis cell area, providing a unitless index between 0 (not plasmolyzed) and 1 (fully plasmolyzed) (Figure S2a). Increasing plasmolysis decreased the percentage of wall-attached plasma membrane in a linear manner (*p* < 0.001). About 70% of the cells overexpressing either LecRK-I.9* or LTI6B exhibited a plasmolysis index between 0.3 and 0.6, indicating that our analysis assessing the density of Hechtian strands was performed within the same range of plasmolysis. This ensured that the observed differences in plasma membrane – wall attachment were not due to variations in plasmolysis levels. However, the slopes presented no statistically significant difference, suggesting that the overexpression of LecRK-I.9* specifically increases the plasma membrane – cell wall connections that form Hechtian strands during plasmolysis (Figure S2a).

As the Hechtian strands are part of a more complex structure(15), we next analyzed the networks of membranes (the Hechtian reticulum) that remained pasted to the wall after plasmolysis at higher spatial resolution. To do so, we extracted the cell surface and segmented the images in three classes: plasma membrane (as viewed from the top), region with Hechtian reticulum, and background. Focusing on the Hechtian reticulum region, we binarized and skeletonized the signal (Figure 2a). Then, we measured the percentage of plasmolyzed area occupied by the Hechtian reticulum at the cell surface. We found that it was significantly higher in cells overexpressing LecRK-I.9* than in LTI6B overexpressing cells, consistent with the observed increase in the density of Hechtian strands. Incidentally, this also demonstrates that such increased density can be independent of plasmodesmata, since the external periclinal cell walls in the epidermis are not in contact with any other cell (Figure 2a).

**Figure 2.**
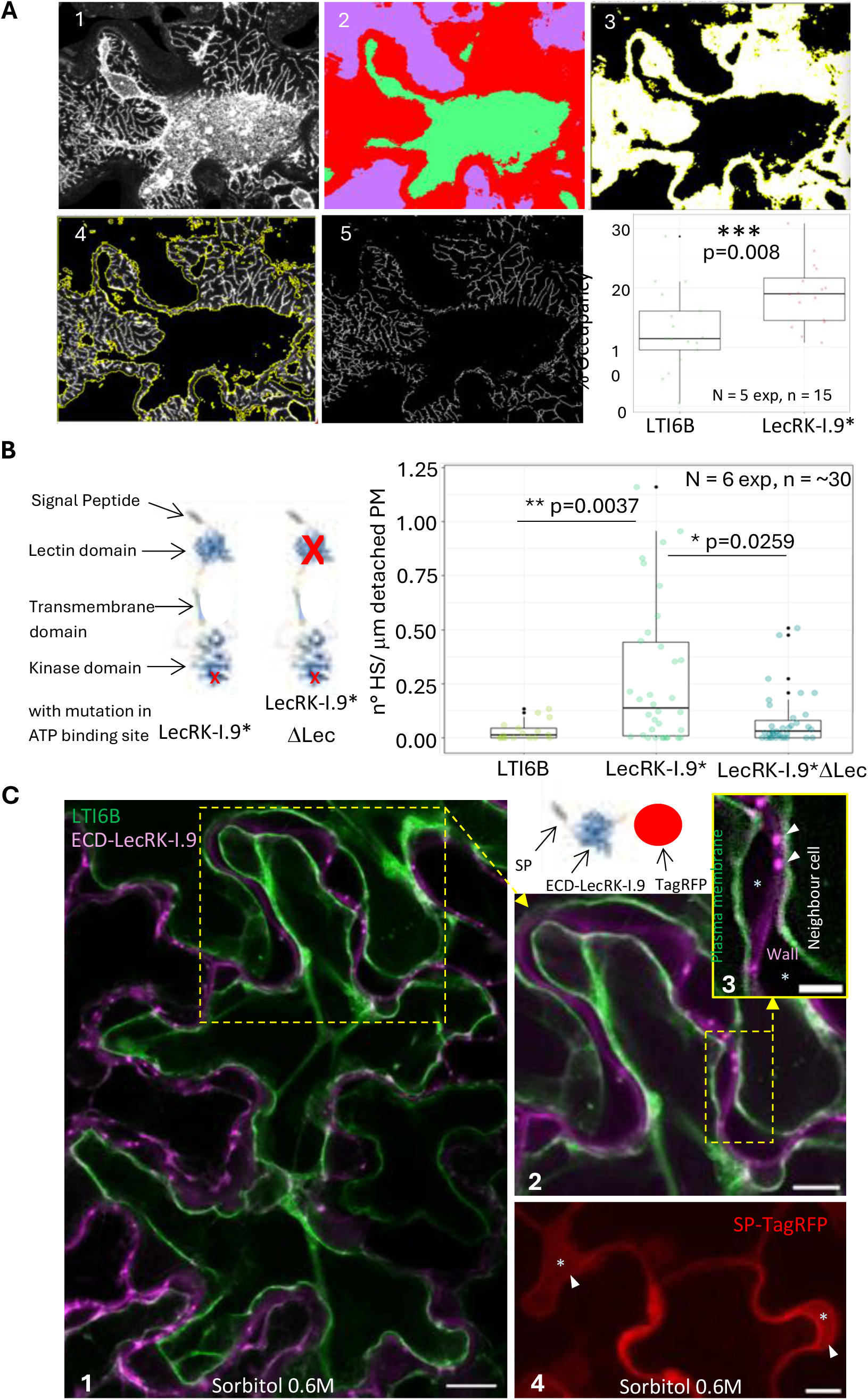
Analysis of LecRK-I.9 contribution to plasma membrane – cell wall connections in *Nicotiana benthamiana*. **A.** Quantification of percentage of the cell surface occupied by the Hechtian reticulum after plasmolysis (% Occupancy). 1. SurfCut projection of the surface. 2. Classified image in plasma membrane (green), area with H. reticulum (red) and background (violet). 3. Mask (black) of plasma membrane and background. 4. Selection (yellow) of area with H. reticulum (white). 5. Binary and skeletonized image. Boxplot representing the % Occupancy for LTI6B and LecRK-I.9* within plasmolyzed areas up to 1000 μm^2^ in size. Statistical significance was determined using the Welch two-sample t-test. Sample size (N = number of experiments, n = number of images with 1 -3 cells averaged) and *p*-value are indicated on the plot. Asterisks denote statistically significant differences compared to the LTI6B control. **B.** Alphafold-predicted structure of LecRKI.9 from Uniprot highlighting the protein domains and the approximate location of the mutations. The boxplot displays the density of Hechtian strands (HS), expressed as number of HS per μm of detached plasma membrane (PM), for LecRK-I.9*, LecRK-I.9*ΔLec and the control LTI6B. Statistical significance was determined using the Kruskal–Wallis test followed by Dunn’s post hoc test with Benjamini–Hochberg correction. No significant difference was observed between LTI6B and LecRK-I.9*ΔLec. Sample size (N = number of experiments, n = number of images with 1 -3 cells averaged) and *p*-values are indicated on the plot. **C.** Transient transformation of *N. benthamiana* followed by plasmolysis of pavement cells with sorbitol 0.6M (30 min.). In green, LTI6B-mCitrine labelling the plasma membrane (and some cytoplasmic streaming). In magenta, the extracellular lectin domain (ECD) of LecRK-I.9 fused to TagRFP (as shown by the scheme), exported to the apoplast by its endogenous signal peptide (SP). In red, the signal of TagRFP exported to the apoplast by the chitinase signal peptide (SP-TagRFP). 1. Entire pavement cell and neighboring cells showing plasma membrane detached from the wall. Scale bar = 15 μm. 2. Inset corresponding to the yellow ROI in 1 showing the signal of ECD-LecRK-I.9 predominantly in the wall. Scale bar = 10 μm. 3. Inset corresponding to the yellow ROI in 2 showing plasmolyzed spaces (*) without ECD-LecRK-I.9 and putative microdomains in the wall that accumulate ECD-LecRK-I.9 (arrowheads). Scale bar = 5 μm. 4. Negative control showing that a protein free in the wall diffuse (arrowheads) into the plasmolized space (*). Scale bar = 10 μm. (Brightness and contrast adjusted in all the images for better observation).

### The lectin domain of LecRK-I.9* is necessary to increase the plasma membrane – cell wall attachment in plasmolyzed pavement cells of *Nicotiana benthamiana*

The extracellular domain of LecRK-I.9 is a L-type lectin, which was described as a carbohydrate binding domain(31). To test whether that lectin domain is required to generate attachment sites, we generated the version of LecRK-I.9* without the lectin domain (LecRK-I.9*ΔLec). Under plasmolysis, the quantification of number of Hechtian strands per μm of detached plasma membrane revealed that the overexpression of LecRK-I.9*ΔLec did not significantly affect the density of Hechtian strands (Figure 2b), when compared to LTI6B in WT cells (*p*-value = 0.2104). To further check this result, we also removed the lectin domain of LecTM and observed a comparable effect when overexpressed (Figure S2b). This supports a scenario in which the lectin domain generates persistent cell wall – plasma membrane attachment sites.

However, there is lack of evidence that the lectin domains of receptor kinases can actually bind the cell wall (32). Hence, we tested the ability of the extracellular lectin domain of LecRK-I.9 (ECD-LecRK-I.9) to remain in the wall during plasmolysis. We reasoned that when the plasma membrane retracts, free proteins in the apoplast would diffuse to the plasmolyzed space while proteins bound to any component of the wall would remain there, in the cell wall. We conducted plasmolysis with sorbitol 0.6M of *Nicotiana benthamiana* pavement cells expressing the membrane marker LTI6B plus the ECD-LecRK-I.9 fused to TagRFP (Figure 2c, Figure S3). We used TagRFP fused to the chitinase signal peptide (SP-TagRFP) as a negative control of a protein exported to the apoplast which cannot bind the wall; and the extracellular domain of RLP4 (ECD-RLP4-TagRFP) as a positive control because it was reported to bind the wall (33) (Figure S3). After plasmolysis, the signal from SP-TagRFP was detected in the plasmolyzed space consistent with the free diffusion of that protein. In contrast, we observed the ECD-LecRK-I.9 signal preferentially located in the wall, with very low to no signal in the plasmolyzed space (Figure 2c, Figure S3). This result supports that the extracellular lectin domain of LecRK-I.9* binds the wall.

### The lectin domain constrains the mobility of LecRK-I.9* clusters in Arabidopsis

To further confirm these results obtained in a heterologous system (*N. Benthamiana*), we next investigated the contribution of the lectin domain of LecRK-I.9 in *A. thaliana*. It has already been reported that LecRK-I.9-TagRFP exhibits very low to no fluorescence when expressed under its native promoter in Arabidopsis (34). As mentioned above, as the overexpression of LecRK-I.9 can be toxic to the plant (Figure S1), we generated a line overexpressing the kinase-dead version of LecRK-I.9 (LecRK-I.9*). Since we are focusing on the structural role of the lectin domain of LecRK-I.9, this is also a way to avoid possible interference with the signaling role of LecRK-I.9. Therefore, to test whether the lectin domain of LecRK-I.9 can generate persistent attachment sites in Arabidopsis, we compared LecRK-I.9* and LecRK-I.9*ΔLec overexpressing lines.

Both LecRK-I.9* and LecRK-I.9*ΔLec versions of the protein localized to plasma membrane, as evidenced by their presence on Hechtian strands similarly to LTI6B (Figure 3a). In water, the distribution of LecRKI.9* and LecRK-I.9*ΔLec appeared largely as homogeneous as LTI6B at the plasma membrane with Airyscan resolution in one single-plane images (Figure S4). We could observe some protein clusters for LTI6B and LecRK-I.9*ΔLec (Figure S4), likely due to transient clustering caused by free lateral diffusion (in addition to overexpression) and steric constraints at the cell cortex (35). In particular, LecRK-I.9*ΔLec exhibited “microtubules corrals” reported before for other plasma membrane proteins (17, 36). After 30 minutes of incubation with sorbitol 0.6M, we observed the formation of large and numerous clusters in both LecRK-I.9* and LecRK-I.9*ΔLec as well as LTI6B cotyledons of 7-d-old overexpressing seedlings (Figure S4).

**Figure 3.**
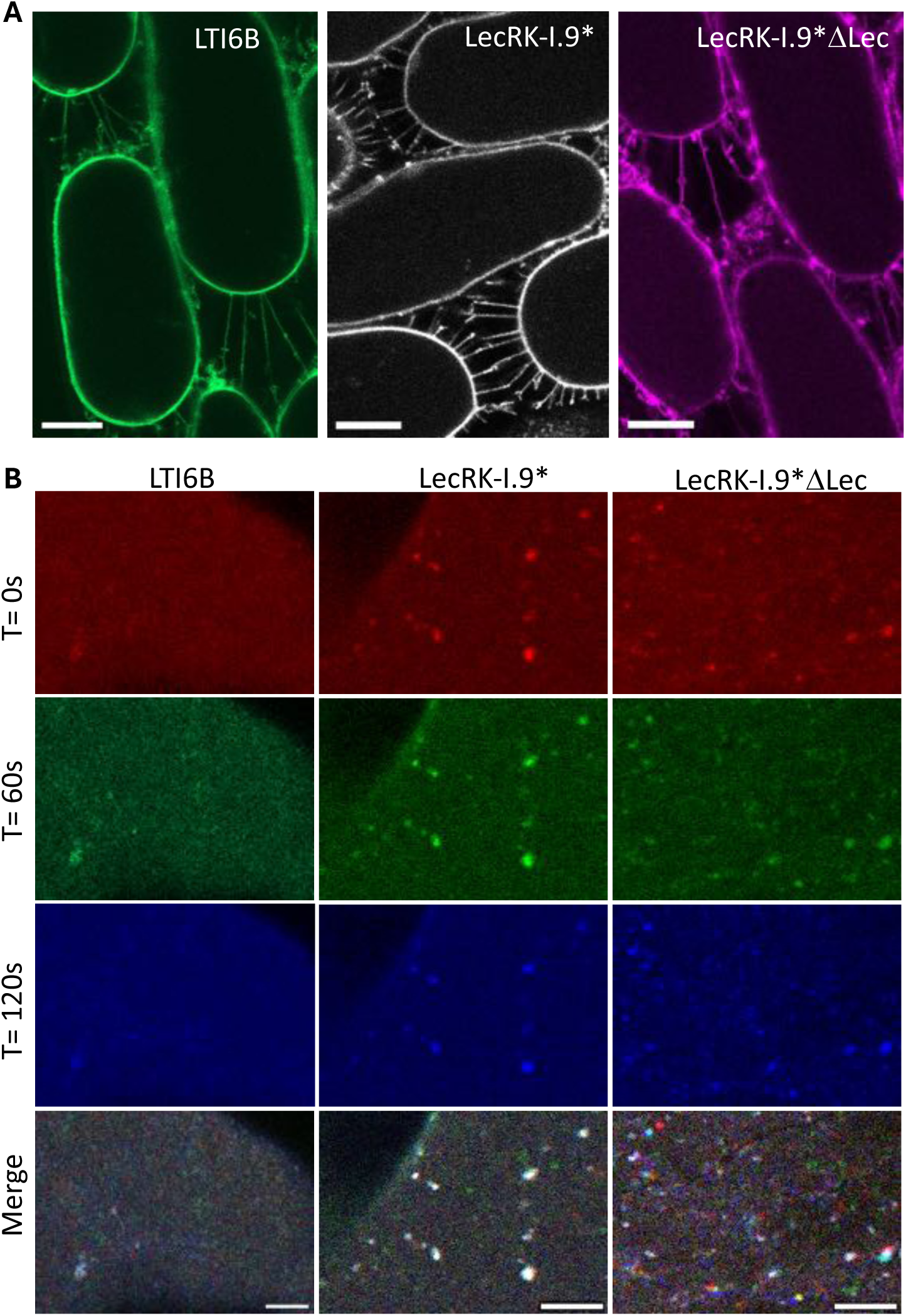
In Arabidopsis, LecRK-I.9* form clusters under hyperosmotic treatments, the mobility of which is lectin dependent. **A.** Confocal images showing LTI6B (green), LecRK-I.9* (grey) and LecRK-I.9*ΔLec (magenta) plasma membrane localization in Hechtian strands of plasmolyzed hypocotyl epidermal cells (at the cell middle-plane) from stably transformed 7 days-old Arabidopsis seedlings (homozygote T3 generation) after sorbitol 0.6M 30-minutes incubation. Scale bar = 10 μm. (Brightness and contrast adjusted in all the images for better observation). **B.** 2-minutes time-lapse series of cotyledon pavement cells surfaces showing the mobility of clusters after incubation in sorbitol 0.6M. Frames are color-coded: Time = 0 seconds (red), Time = 60 seconds (green), and Time = 120 seconds (blue). Clusters that did not move appear white in the merge images. Scale bar = 5 μm. (Brightness and contrast adjusted in all the images for better observation).

The passive formation of protein clusters can be the consequence of changes in membrane topology and biophysical properties (e.g. tension) provoked by hyperosmotic stress (37, 38); that may be amplified by aggregation due to overexpression (39). Here we took advantage of this phenomenon to analyze the contribution of the lectin domain in the mobility of clusters. Two-minute-time-lapse imaging of Arabidopsis cotyledon pavement cells incubated with sorbitol 0.6M, revealed that there are populations of clusters with different mobility in all cases. However, the mean displacement of LecRK-I.9* clusters during 2 minutes was 0.19 ± 0.18 μm (SD) with an average speed of 0.04 ± 0.06 μm/s (SD); while the mean displacement of LecRK-I.9*ΔLec clusters in that time range significantly increased to 0.55 ± 0.64 μm (*p*-value = 6.081^e-08^) with an average speed of 0.14 ± 0.14 μm/s (*p*-value = 0.00341). This means that the majority of LecRK-I.9* clusters presented a mobility close to zero, in contrast to highly mobile LecRK-I.9*ΔLec clusters (Figure 3b). Another way to see this, is revealing movement in one image by merging the frames at times 0, 1 and 2 minutes. Strikingly, most of LecRK-I.9* clusters did not move in that time range (clusters that appeared in white), while most of LecRK-I.9*ΔLec clusters appeared in different colors and positions on the merge images revealing their movement (Figure 3b). A similar response was obtained at milder hyperosmotic treatment (sorbitol 0.3M, Figure S5). These results indicate that the lectin domain reduces the mobility of LecRK-I.9*, most likely due to anchoring to the wall; with the formation of nearly immobile clusters that can putatively mark plasma membrane – cell wall attachment sites.

### LecRK-I.9* cluster distribution correlates with predicted differences in wall properties, only when the lectin domain is present

If LecRK-I.9* cluster immobility reflects their attachment to the wall, we reasoned that different wall properties or compositions may also alter cluster density. To test this hypothesis, we analyzed the distribution of LecRK-I.9* and LecRK-I.9*ΔLec clusters at the plasma membrane under hyperosmotic stress (30 min with 0.6M sorbitol) in cotyledons and hypocotyls; comparing the outer and inner walls of the epidermis. Indeed, in cotyledons, the outer face of the epidermis is thicker and mechanically reinforced, because it is more load-bearing than the inner face of the epidermis, as shown for instance with the microtubule response to stress (40). In contrast, computational modeling together with experiments rather suggest that the load-bearing face in hypocotyls is on the inner side of the epidermis (41–44). Therefore, we segmented the clusters and analyzed their density (number of clusters per μm^2^ of plasma membrane area) and size (area as viewed from the top, in μm^2^) on both the outer and inner surfaces of epidermal cells in cotyledons and hypocotyls (Figure 4, Figure S6a). Note that despite transient exposure to harsh hyperosmotic stress, the cells we analyzed were able to maintain most of their volume (Figure S6b) (likely due to the high osmoregulation capacity of epidermis, (45)). Note also that seedlings could survive at least 5 days after recovery 1h in water from the harsh hyperosmotic treatment we performed (Figure S6c).

**Figure 4.**
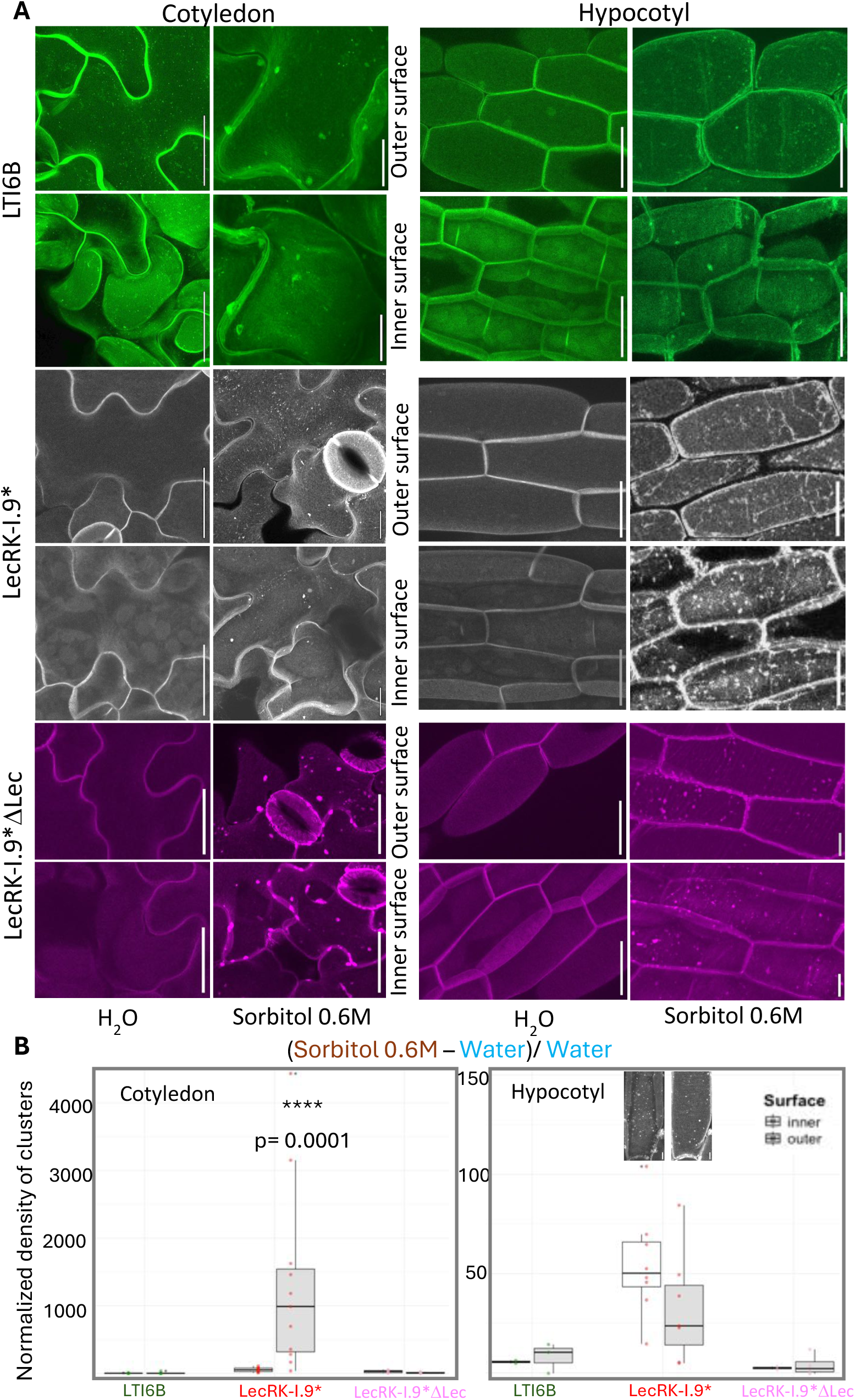
LecRK-I.9* cluster distribution correlates with predicted differences in wall properties, only when the lectin domain is present. **A.** Confocal images showing outer and inner surfaces of cotyledon and hypocotyl epidermal cells from 7-days-old seedlings incubated in water (control condition) or sorbitol 0.6M during 30 minutes (hyperosmotic condition). The patterns of clusters are shown for LTI6B (in green), LecRK-I.9* (in grey) and LecRK-I.9*ΔLec (in magenta). Scale bars = 25 μm; except for LecRK-I.9* and LTI6B cotyledons in sorbitol and hypocotyls of LecRK-I.9*ΔLec in sorbitol where scales bars = 10 μm. All images of LecRK-I.9* and LTI6B cotyledons were acquired with Airyscan detector. (Brightness and contrast adjusted in all the images for better observation). **B.** Boxplots presenting the number density of clusters for LTI6B, LecRK-I.9* and LecRK-I.9*ΔLec in the hyperosmotic condition normalized by the control condition ((Sorbitol 0.6M – Water)/ Water). Inner vs. Outer surfaces within each genotype were compared by Wilcoxon test. N = 3 experiments, n = ∼3-9 plants/genotype/treatment (average 1-3 cells per image). Asterisks denote significant differences. *p*-value is indicated on the cotyledon plot. Images on the hypocotyl plot highlight the strong trend of asymmetrical pattern displayed by LecRK-I.9* clusters in the inner and outer surface of the hypocotyl cell. Scale bar = 10 μm.

Following treatment with 0.6M sorbitol, the density of LecRK-I.9* clusters significantly increased on both surfaces and in both organs compared to water-treated controls, as expected (Figure 4a, Figure S6a). Interestingly, this increase was accompanied by a marked asymmetry in the distribution of clusters between inner and outer surfaces of cotyledons and hypocotyls: LecRK-I.9* clusters were preferentially located on the outer surface of cotyledons and inner surface of hypocotyls epidermal cells (Figure 4a,b; Figure S6a). Such asymmetry was not observed for LecRK-I.9*ΔLec or LTI6B cluster localization (Figure 4a,b; Figure S6a). Thus, the density of LecRK-I.9* clusters positively correlate with predicted mechanical reinforcement of the wall, in a lectin domain dependent way. This confirms that the lectin domain is required for attachment to the wall, and suggests that its attachment may also reflect wall properties.

### LecRK-I.9* overexpressing lines can expand their cotyledons and leaves despite the low water potential conditions

Both the reproducible formation of Hechtian strands under plasmolysis, and the preferential attachment of the lectin domain to certain walls suggest that persistent cell wall - plasma membrane attachment sites may have physiological relevance. Hence, we tested whether the promotion of attachment sites in LecRK-I.9* overexpressors would impact the plant response to low water availability conditions.

To do so, we transferred 5-d-old seedlings grown on normal medium (0.07 ± 0.02 Osmol/L) to PEG-infused plates (0.18 ± 0.02 Osmol/L) in which the water potential is reduced. In parallel, we also transferred 5-d-old seedlings grown on normal medium to fresh control medium at the same osmolarity (0.07 ± 0.02 Osmol/L) (Figure 5). After 10 days in the treatment, we assessed the ability of the plants to continue growing despite the low availability of water (Figure 5a). We considered a plant as resistant if it was able to expand the cotyledons and leaves achieving a rosette architecture comparable to the one in mock conditions (Figure 5b, Figure S7). Despite the high variability of such physiological assays, LecRK-I.9* overexpressing lines exhibited significantly higher frequency of resistant plants. On average 57 ± 19 % (mean ± standard deviation) of LecRK-I.9* seedlings were able to expand their cotyledons and leaves, while 29 ± 22 % of WT seedlings were able to do so (\*\*\**p*-value = 0.003). Strikingly, LecRK-I.9*ΔLec seedlings were as sensitive as WT exhibiting a frequency of 21 ± 20 % for resistant plants (*p*-value = 0.519), and significantly different from LecRK-I.9* (\*\*\**p*-value = 0.002, Figure 5c). Therefore, the promotion of cell wall - plasma membrane attachment site through the lectin domain of LecRK-I.9* can confer resistance to low water potential conditions.

**Figure 5.**
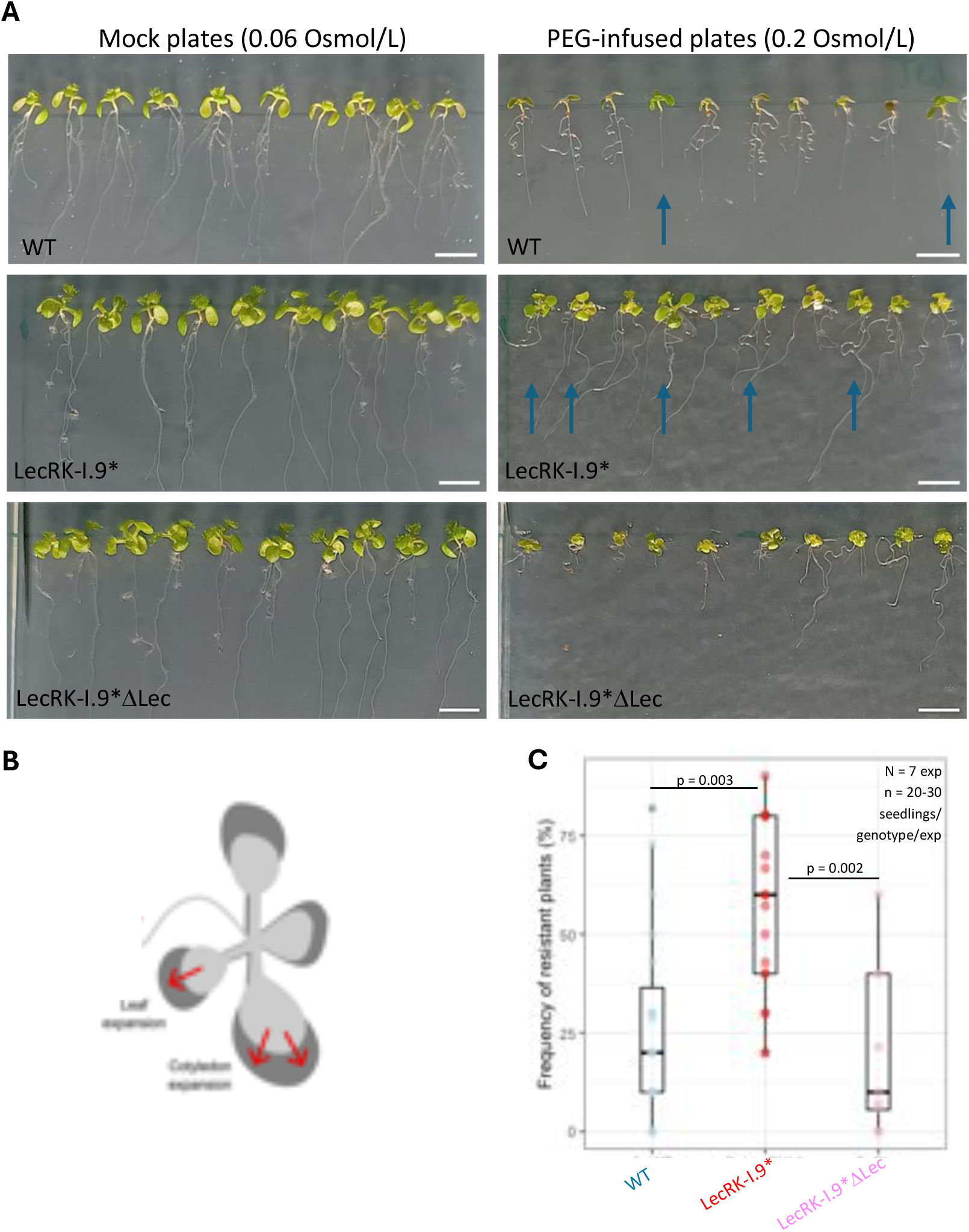
LecRK-I.9* overexpressing lines can expand their cotyledons and leaves despite the low water potential stress. **A.** Images showing WT (Col-0) Arabidopsis seedlings, LecRK-I.9* and LecRK-I.9*ΔLec overexpressing lines after 10 days from being transferred to mock plates (0.06 Osmol/L) or PEG-infiltrated plates (0.2 Osmol/L). See Material and methods for details about the assay and plate preparation. Blue arrows show examples of resistant plants. Scale bars = 5mm. **B.** Scheme depicting the criteria for considering a plant as resistant: expansion of cotyledons and leaves achieving a rosette architecture comparable to the one in mock conditions. **C.** Boxplot showing the frequency of resistant plants for each genotype across 7 experiments (N), each one with 2 to 3 plates per treatment (Mock/ PEG) containing 7 -10 seedlings of each genotype per plate (n = 20 – 30 seedlings per genotype per experiment). Statistical significance was determined using the Kruskal–Wallis test followed by Dunn’s post hoc test with Benjamini–Hochberg correction. Sample size and *p*-values are indicated on the plot.

## DISCUSSION

After plasmolysis, we observed that the plasma membrane – cell wall continuum is structurally constrained at discrete sites in *Arabidopsis thaliana* and *Nicotiana benthamiana*, as was reported before for several species (15). We found that the overexpression of LecRK-I.9* not only increases the density of Hechtian strands (Figure 1e), but it also increases the density of the Hechtian reticulum (Figure 2a), thus showing that the effect can be independent of the density of plasmodesmata. This provides the first clue in our quest for a molecular component able to mediate attachment between the cell wall and the plasma membrane, leading to the formation of Hechtian strands upon plasmolysis

To formally demonstrate that the attachment requires the lectin domain, we showed that (i) the free lectin domain (ECD-LecRK-I.9) remain localized to the cell wall during plasmolysis (Figure 2c, Figure S3), (ii) LecRK-I-9* clusters are largely immobile (Figure 3b, Figure S5), (iii) that the deletion of the lectin domain in LecRK-I-9*ΔLec leads to WT density of Hechtian strands (Figure 2b) and mobile clusters (Figure 3b, Figure S5). Thus, we propose that the lectin domain of LecRK-I.9 can generate persistent connections between the plasma membrane and the cell wall, with implications for Hechtian strand density.

LecRK-I.9 belongs to the legume-type lectin protein kinase family (19, 46). Legume-type (L-type) lectins form a diverse family of carbohydrate-binding proteins sharing significant similarities in their structures, but remarkable variability in their carbohydrate-binding specificities (31). There are few examples of extracellular domains of receptor kinases that can bind specific components in plant cell walls. For instance, the MalectinA domain of FERONIA interacts with unesterified pectin (47), and the extracellular domain of the wall-associated kinase WAK1 can bind pectin such as polygalacturonic acid oligogalacturonides (48). However, further research will need to explore the interaction between the legume-type lectin of LecRKs and specific targets in the cell wall (32). Note that the overexpression of LecGPI, which also has a legume-type lectin domain, did not increase the density of Hechtian strands (Figure 1e). This means that the lectin domain is necessary but not sufficient for adhesion: a GPI anchor might be too weak of an anchor (49), when compared to a receptor with a transmembrane domain, if the protein is not interacting with any other protein that may function as a linker. Consistently, the overexpression of COBRA, another GPI anchor protein involved in anisotropic root cell expansion(24), did not promote Hechtian strands formation (Figure 1e). Conversely, AGP18 despite of having a GPI anchor, had a mild effect on increasing density of Hechtian strands (Figure 1e). This effect may be explained by the reported interaction between AGP18 and the receptor kinase CERK1 confirmed by yeast two hybrid assay (50).

Our results also suggest that the attachment of the lectin domain to the wall may depend on wall properties and composition. In particular, we observed that LecRK-I.9* clusters are enriched on the cell faces that are predicted to be mechanically reinforced before plasmolysis (40) (the outer epidermal wall of the cotyledon and the inner epidermal wall of the hypocotyl). Again, this asymmetry is abolished when removing the lectin domain (Figure 4, Figure S6a). This is consistent with the role of the lectin domain at cell wall – plasma membrane attachment sites, but it also suggests that the lectin domain, and the related density of LecRK-I.9* clusters, may reflect the wall properties. Recently, it was observed that treatments with 1 M sorbitol immobilized CSC (Cellulose Synthase Complex) into nanodomains at the plasma membrane, which showed colocalization with GFP-LTI6B-labeled membrane-wall attachment sites in Arabidopsis roots (7). This highlight that clusters of a plasma membrane protein complex that impacts the composition of the cell wall, can pattern plasma membrane – wall attachment sites during hyperosmotic stress. How the mechanical reinforcement of the wall affects the structure and the signaling of protein complexes at the cell cortex is an open question. For instance, the LRX8-RALF4-pectin and the LRX1/2-RALF22-pectin interactions, lead to the compaction of pectin and these complexes become integral part of the cell wall (51, 52). The presence of persistent attachment sites through the lectin domain may help to generate static hubs, which could help the cell to sense relative deformations in the wall.

It has been suggested that Hechtian strands might be important for resisting conditions that challenge the integrity of the plasma membrane – cell wall continuum (7, 53), such as intermittent osmotic stress derived by cold, drought, salinity (54, 55), as well as wounding or pathogen infection (56); and mechanical stress derived by development (20). Here we find that the overexpression of LecRK-I.9* confers seedling resistance to low water potential conditions (Figure 5, Figure S7). Importantly, this effect is cancelled by the deletion of the lectin domain. This provides an important building block for the further exploration of the link between plasma membrane – cell wall attachment and plant resistance to water stress. Recently, FERONIA was characterized as an osmosensor, capable of sensing turgor reduction and consequently triggering nanodomain formation and kinase activation (57). In that study, the cell wall anchoring mediated by the extracellular MalectinA domain, was essential for initiating the osmotic signaling cascade that regulates root hydrotropism response and water loss (57). In this context, our results provide evidence of another extracellular carbohydrate binding domain of a receptor kinase, specifically involved in Hechtian attachment sites, that influences the clustering dynamics of the protein during osmotic stress. The persistent attachment of the plasma membrane to the wall thus emerges as physiologically relevant for plants during low water potential conditions in which there is less availability of water.

## MATERIALS AND METHODS

### Plant material and growth conditions

All the experiments were performed using either *Nicotiana benthamiana* wild-type plants, transiently transformed with the constructs specified in the Table S1; or *Arabidopsis thaliana* ecotype Col-0 plants, expressing the following constructs:

- LTI6B: (AT3G05890) *p35S::eGFP-LTI6B*(*58, 59*), used as a membrane marker and control in Arabidopsis.
- LecRK-I.9 and LecRK-I.9* (*p35S::LecRK-I.9-TagRFP* and *p35S::LecRK-I.9*-TagRFP*, respectively), lines overexpressing the full-length LecRK-I.9 (At5g60300) protein fused to the Red Fluorescent Protein. LecRK-I.9 corresponds to the wild-type version and LecRK-I.9* carries 5 alanine substitutions at positions 350aa – 354aa within the ATP binding motif LGKGG.
- LecRK-I.9*ΔLec (*p35S::LecRK-I.9*ΔLec-TagRFP*), a line that overexpresses a truncated form of LecRK-I.9* in which its extracellular lectin was removed (D244aa at positions 22aa – 265aa).

Arabidopsis seeds were surface-sterilized with chlorine gas (4ml HCl2 37% in 100mL sodium hypochlorite) for 4 hours and sown in petri dishes containing half-strength Murashige and Skoog (1/2 MS) medium supplemented with vitamins (Sigma-Aldrich, St Louis, MO, USA) and 0,8% Agar, at pH 5.7 (KOH). After 3 days of stratification at 4°C in the dark, the plates were placed in the culture chambers under a 16h light/ 8h dark cycle within a 24-hour photoperiod at 22°C during 7 days, or the number of days specified for each assay.

### Cloning and generation of Arabidopsis transgenic lines

All the constructs specified in the Table S1 were generated using the Gateway® cloning technology. The genes were amplified by PCR from the *A. thaliana* Col-0 genomic DNA using AccuPrime™ Taq DNA Polymerase High Fidelity (Invitrogen®, Carlsbad, CA, USA) and the primers detailed in Table S1; except for LecRK-I.9 which was amplified from the vector pDA10051 carrying the complete coding sequence of AT5G60300, referenced in the databases of the RIKEN research center and described on the website https://www.brc.riken.jp/lab/epd/catalog/cdnaclone.html.

All the PCR fragments were cloned in the *pDONR207* (ampicillin 100 µg/mL) entry vector by BP reactions (Invitrogen®), transformed in *E. coli* TOP10 (Invitrogen) and sequenced (Eurofins Genomics). Expression vectors were generated by LR reactions using the *pDONR207* entry clones and the destination vector *pEAQ-HT-DEST1* (C. Ritzenthaler IBMP Strasbourg) that contains the *P19* gene under the control of a *35S* promoter, which is a suppressor of the RNA interference silencing to promote the accumulation of heterologous proteins in *N. benthamiana* transient expression. The constructs for AGP18, LecGPI and COBRA were cloned without their native signal peptides (which are cleaved upon secretion to apoplast) into *pEAQ-HT-DEST1* vectors containing the signal peptide (SP) of Chitinase fused to TagRFP, resulting in fusion proteins of the form SP-TagRFP(N-terminal)-AGP18/ LecGPI/ COBRA. This design ensured proper secretion while avoiding disruption of the C-terminal GPI anchor, which would be incompatible with a C-terminal fluorescent tag. The *Agrobacterium tumefaciens* GV3101::pMP90 strain was transformed with the recombinant vectors and transformant colonies were selected by antibiotic resistance (gentamycin 10 µg/mL, kanamycin 50 µg/mL, rifampicin 50 µg/mL).

Site-directed mutagenesis was carried out using the ’Q5® Site-Directed Mutagenesis’ kit (New England BioLabs) with the primers specified in Table S1 which were designed using the supplier’s software (NEBaseChanger.neb.com). Site-directed mutagenesis was implemented to substitute the 15bp: CTGGGAAAAGGAGGT (positions 1048bp – 1062bp) by GCTGCAGCAGCTGCA of *LecRK-I.9* gene in *pDONR207* to generate LecRK-I.9*; and to delete the lectin domain of LecRK-I.9* and LecTM to generate DLec (D732bp = positions 64bp – 795bp) and LecTMDLec (D714bp = positions 82bp - 795bp) respectively. Site-directed mutagenesis was also implemented to delete *P19* from the destination vector to stably transform Arabidopsis plants with the floral dip method(60). Homozygous T3 progenies were used for all experiments except for the ones in Figure S1 where T2 generation was used.

### Screening and plasmolysis assays in *N.benthamiana*

Wild-type *Nicotiana benthamiana* plants were used for transient transformations by agroinfiltration. The plants were grown in a growth chamber for 4 weeks with a photoperiod of 16 hours of light / 8 hours of darkness at a temperature of [25/22]°C and a humidity of 70%. The transformed agrobacteria with the constructs specified in the Table S1 were cultured overnight in liquid LB medium at 28°C. After centrifugation, the bacterial pellets were resuspended in a solution of 10 mM MgCl₂ and 150 µM acetosyringone to obtain an OD₆₀₀nm of 0.5, then kept at room temperature for 4 hours. Leaves from positions 3, 4, and 5 of *N. benthamiana* plants were infiltrated using a 1 mL syringe (without needle) on the abaxial face of the leaves. After 48 and 72 hours from infiltration, leaf discs were collected using a hole punch and placed into a syringe with glycerol 10% to be degassed and infiltrated with the osmoticum. The procedure was repeated with calcofluor (Fluorescent brightener 28, Sigma-Aldrich) 0,1% (w/v) to stain the cell wall. The leaf discs were incubated for 20 minutes in glycerol10% in order to plasmolyze the epidermal cells (and reach a steady state) before imaging. For each experiment, 3 leaf discs per protein candidate were imaged in several sectors during 40 minutes, with random spatial sampling and avoiding the borders. *pPDF1::LTI6B-mCitrine* was used as an epidermal membrane marker and control (*pBART* vector, spectinomycin 100 µg/mL; co-infiltrated with P19). Each experiment was repeated for a minimum of 3 times. Same procedure was followed to image *N. benthamiana* pavement cells transiently transformed with *pPDF1::LTI6B-mCitrine* plus *p35s::ECD-LecRK-I.9-TagRFP/ pUBQ10::ECD-RLP4-TagRFP/ p35S::SP-TagRFP* but the osmoticum used for plasmolysis was sorbitol 0.6M.

### Confocal microscopy imaging

Confocal images of leaf discs were acquired using a spectral laser scanning confocal system (SP8, Leica) equipped with a microscope (DMi8, Leica), using a 25x water immersion objective (Fluotar VIZIR N.A. 0.95). Images of 1024 x 1024 px or 512 x 512 px were acquired in sequential mode averaging twice the signal per frame. The configuration for each fluorophore was: LTI6B-mCitrine (λ excitation: 488nm, λ emission: 500–540nm, detector gain: 700V, laser: 30%); Calcofluor (λ excitation: 405nm, λ emission: 430–475nm, detector gain: 400V, laser: 5 %): candidate-TagRFP (λ excitation: 561nm, λ emission: 576–628nm, detector gain: 500V, laser: 15 %). z-Stack series (voxel size 0.2 µm x 0.2 µm x 1 µm) were acquired from plasmolyzed pavement cells in the abaxial epidermis.

Arabidopsis seedlings were incubated for 30 minutes (to reach a steady state for the working time frame) in water (control condition) or sorbitol 0.6M/ 0.3M (conditions mostly used as high osmotic stress(61)) before imaging with a Confocal microscope Zeiss LSM980 using a 40x/NA 1.20 water immersion objective and GaAsP detector. Images of 930 x 930 px were acquired with excitation and emission settings optimized for each fluorophore, averaging twice the signal per line. The configuration for each fluorophore was: eGFP-LTI6B (λ excitation : 488 nm and λ emission: 499-588 nm; Laser Power / Detector Gain: for cotyledons 0.3%-0.4% / 800V; for hypocotyls 0.5-0.8% / 850V) ; LecRK-I.9*-TagRFP (λ excitation: 561 nm and λ emission: 570-659 nm; Laser Power / Detector Gain: 0.2-0.5%/ 850V); LecRK-I.9*ΔLec -TagRFP (λ excitation: 561 nm and λ emission: 570-659 nm; Laser Power / Detector Gain: 0.1%-0.3% / 800V). The autofluorescence of chlorophyll was collected between λ 683 and 720 nm (Detector Gain: 650V). z-Stack series (voxel size 0.1 µm x 0.1 µm x 0.5 µm) were acquired from epidermal cells of cotyledons and hypocotyls for each genotype in the two different conditions. For each experiment, two to three plants per genotype x organ x treatment were imaged and each experiment was replicated at least 3 times. High-resolution imaging was performed using the Airyscan detector. Airyscan acquisition was conducted in super-resolution mode, and raw data were processed using Zeiss Zen software with default deconvolution settings. For time-lapse acquisition of one single plane at the cell surface, cotyledons were mounted on slides containing a low toxicity silicone adhesive (KWIK-SIL, WPI, USA) after the hyperosmotic treatment. 2-minutes time series were acquired with no interval.

## QUANTIFICATION AND STATISTICAL ANALYSIS

### Quantification of Hechtian Strands

We developed a simple pipeline to quantify the number of Hechtian strands per µm of detached plasma membrane (density), available in https://github.com/Denise-arico/PlasmaMembrane_wall_attachment_quantification. Images of plasmolyzed pavement cells of *N. benthamiana* in the LTI6B-mCitrine channel (for control and candidates verifying they had signal on the TagRFPchannel) were processed in Fiji (62) (https://imagej.net/software/fiji/, Version: 2.16.0/1.54p) to obtain constant 10 µm-depth z-stack maximal projections avoiding the cell surface (with the Hechtian reticulum). The selection of Regions Of Interest (ROI) were performed manually, as lines parallel to the detached plasma membrane and perpendicular to the Hechtian strands (Figure 1d). The fluorescence intensities vs. the lengths of the ROIs were plotted, and the number of picks (Hechtian strands) with intensity value higher than a threshold of 40 were counted (Figure 1d). This procedure was done for every plasmolytic space of all the neighboring cells with signal, obtaining an average value for each image.

### Quantification of Hechtian Reticulum

We developed a pipeline to quantify the percentage of the cell surface occupied by the Hechtian reticulum after plasmolysis (% Occupancy), available in https://github.com/Denise-arico/PlasmaMembrane_wall_attachment_quantification. Surfaces of plasmolyzed *N. benthamiana* pavement cells were extracted using SurfCut(63), making a projection of the first 5 slices with a manual threshold value and radius of 20 (Figure 2a-1). The images were segmented in Fiji (Version: 2.16.0/1.54p) using the Trainable Weka Segmentation tool(64) in 3 classes: Plasma Membrane, Plasmolyzed area with Hechtian reticulum and Background (Figure 2a-2). Plasma membrane and background were masked on the segmented images and the plasmolyzed area with Hechtian reticulum was selected (Figure 2a-3) in order to restore that selection on the original SurfCut projection image where the plasmolyzed area was measured (Figure 2a-4), binarized and skeletonized (Figure 2a-5). The percentage of occupancy was calculated as the number of pixels with value 255 (Hechtian reticulum) divided by the total number of pixels in the plasmolyzed area, all multiplied by 100. Three cells were analyzed and averaged per image.

### Quantification of degree of plasmolysis and percentage of plasma membrane attached to the wall

We developed a pipeline, available in https://github.com/Denise-arico/PlasmaMembrane_wall_attachment_quantification, to quantify the cell degree of plasmolysis and the percentage of plasma membrane that remained attached to the wall. Middle-plane of plasmolyzed *N. benthamiana* pavement cells were analyzed in Fiji (Version: 2.16.0/1.54p). The calcofluor channel was used to segment the individual cells and measure their perimeter and area (Figure S2a-1) as an approximation of total plasma membrane length and area before plasmolysis, respectively. The area of the shrunken protoplast of the segmented cells was measured as the area after plasmolysis (Figure S2a-2). The selection of ROIs was performed manually where segments of plasma membrane remained attached to the wall (Figure S2a-3). The degree of plasmolysis for each cell was calculated as the difference of area before and after plasmolysis divided by the area before plasmolysis (plasmolysis index). The percentage of plasma membrane attached to the wall was calculated by the sum of ROIs lengths divided by the total plasma membrane length. This procedure was repeated for 1 to 3 cells so as to obtain an average value per image.

### Quantification of clusters in Arabidopsis

For each genotype, organ and condition, the inner and outer surfaces of the cells were extracted by maximum intensity z-stack projections in Fiji (Version: 2.16.0/1.54p). The resulting images were segmented in 3 classes by pixel classification using the trained-classifier ilastik(65) ^(^www.ilastik.org, 1.4.1rc2-arm64-OSX): Plasma membrane, Clusters and Background (Figure S6a1-3). The segmented images were binarized in Fiji, where the clusters were set to white (pixel value 255 for an 8-bit image) and the background and plasma membrane were masked (pixel value 0 or black). The selection of ROIs (area of cell without borders) was performed manually on the original maximal projection images (Figure S6a4). The area of the cell and the density of clusters (the number of clusters per area of cell) were computed automatically in Fiji, for 2 - 5 cells per image, with a macro script developed by Vincent Bayle (RDP, ENS-Lyon, France), available in https://github.com/RDP-vbayle/SiCE_FIJI_Macro/blob/main/misc/MAcroDots%20Arico%20et%20al.2026%20250210.ijm. The mobility of the clusters was quantified using the plug-in TrackMate (66) in Fiji.

### Statistical Analysis

All plots and statistical analysis were performed in R (https://www.R-project.org/, Version 2024.12.1+563). Normality and homoscedasticity were tested for each dataset in each experiment using the Shapiro-Wilk test and Levene’s test respectively. In all the cases normality was rejected, so comparisons were done using the non-parametric Kruskal-Wallis test, followed by Dunn’s post-hoc test with Benjamini-Hochberg correction for multiple comparisons; or Wilcoxon rank-sum test in cases of two- groups comparisons. Details regarding number of samples and significance are specified in the figure legends.

To investigate how the percentage of plasma membrane attached to the wall varied with the degree of plasmolysis, we performed a linear regression in R (Version 2024.12.1+563) using the lm() function. The model included the degree of plasmolysis as a continuous variable representing the severity of plasmolysis), genotype (a categorical factor with levels “LTI6B” and “LecRK-I.9*”), and their interaction as predictors of the percentage of plasma membrane attached.

### Physiological assays with PEG-infused plates

*Arabidopsis thaliana* seedlings (WT Col-0 and the overexpression lines LecRK-I.9* and LecRK-I.9*ΔLec; see Plant Material section) were grown vertically for 5 days under long-day conditions (16 h light, 22°C, 100 µmol m⁻² s⁻¹) on 0.5× MS agar medium (0.8% agar, pH 5.7). Seedlings were then transferred to polyethylene glycol (PEG)-infused or mock plates.

PEG solution was prepared by dissolving 8 g of PEG 20000 in 40 mL of 0.5× MS liquid medium (pH 5.7) supplemented with 40 mL of Plant Preservative Mixture (PPM; 1:1000, Plant Cell Technology). The solution was poured onto MS agar plates and incubated overnight to allow diffusion and equilibration of water potential. Excess solution was removed the next day. Osmolarity decreased from 0.6 Osmol L⁻¹ ± 0.05% (initial solution; measured with an Osmomat 030 cryoscopic osmometer) to 0.2 Osmol L⁻¹ after equilibration. Mock plates were prepared similarly using MS liquid medium with PPM only; osmolarity remained at 0.06 Osmol L⁻¹ before and after incubation. This protocol was adapted from (67–69).

After 10 days on mock or PEG-infiltrated plates, seedling resistance to low water availability was assessed. Seedlings were considered resistant if they exhibited substantial cotyledon and leaf expansion, forming a rosette architecture comparable to that observed under mock conditions. Each experiment included 2–3 PEG plates and an equal number of mock plates, each containing 7–10 seedlings per genotype, with randomized positioning. For each plate, frequency of resistant plants (%) was calculated as the number of resistant seedlings divided by the total number of seedlings per genotype, and values were averaged per experiment. Statistical analyses are described in the figure legends. Seedlings lacking significant/ clearly visible cotyledon and leaf expansion were classified as non-resistant in all cases.

## ABBREVIATIONS

LTI6B: Low Temperature-Induced Protein 6B.
LecRK-I.9*: Legume-type lectin Receptor-like kinase I.9 (also called DORN1 or P2K1). * 5 alanine substitutions at positions 350aa – 354aa within the ATP binding motif.
LecRK-I.9*ΔLec: LecRK-I.9* without lectin domain (Δ244aa at positions 22aa – 265aa).

## ACKNOWLEDGEMENTS

We thank the Cell Wall team at the LRSV laboratory in Toulouse, the MechanoDevo team, and all members of the RDP laboratory in Lyon for their insightful discussions. In particular, we are grateful to Vincent Bayle (RDP, ENS-Lyon), Aurélie Le Ru (FRAIB microscopy platform, Toulouse), and Adelin Barbacci (LIPME, Toulouse) for their collaboration in designing the imaging quantification; and to Fahd Rachad for generating the first clone of LecRK-I.9 during his Master in the LRSV, Toulouse.

## FUNDING

This work was supported by the European Research Council (ERC-2021-AdG-101019515, “Musix”).

## RESOURCE AVAILABILITY

### Lead Contact

Request for further information and resources should be directed to, and will be fulfilled by the lead contact, Denise S. Arico.

### Materials availability

All reagents generated in this study are available from the lead contact without restriction.

### Data sharing plan

- All original codes for quantification used in this study have been deposited to GitHub and are publicly available at https://github.com/Denise-arico/PlasmaMembrane_wall_attachment_quantification.
- Any additional information required to reanalyze the data reported in this paper is available from the lead contact upon request.

## Notes

### Competing Interest Statement

The authors have declared no competing interest.

### Summary of Updates

- New data on extracellular domain (lectin domain) in the cell wall after plasmolysis - New data on LecRK-I.9 overexpressor resistance to water stress - Focus on the lectin domain (comparison between LecRK-I.9 overexpressor and LecRK-I.9 DeltaLec overexpressor)

