## Supporting Information for "A molecular glue in plants: The lectin domain of LecRK-I.9 creates persistent plasma membrane – cell wall connections"

**This PDF file contains**

Graphical abstract

Figures S1 – S7

Table S1

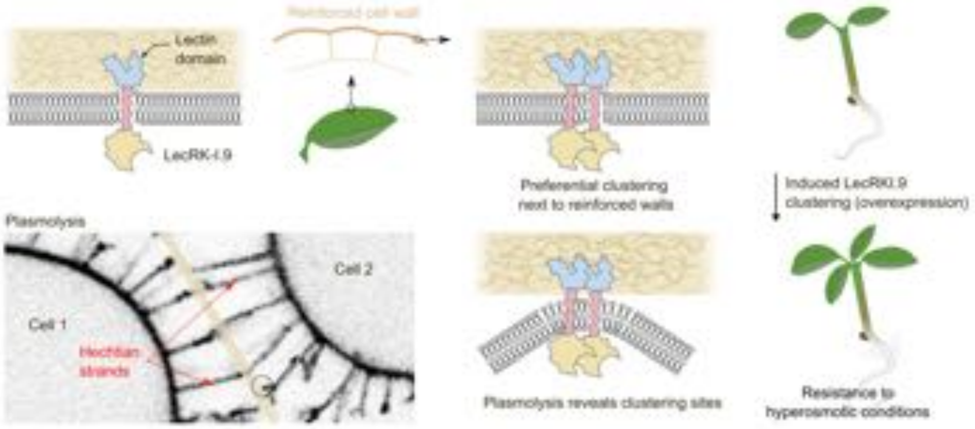


**Graphical abstract**

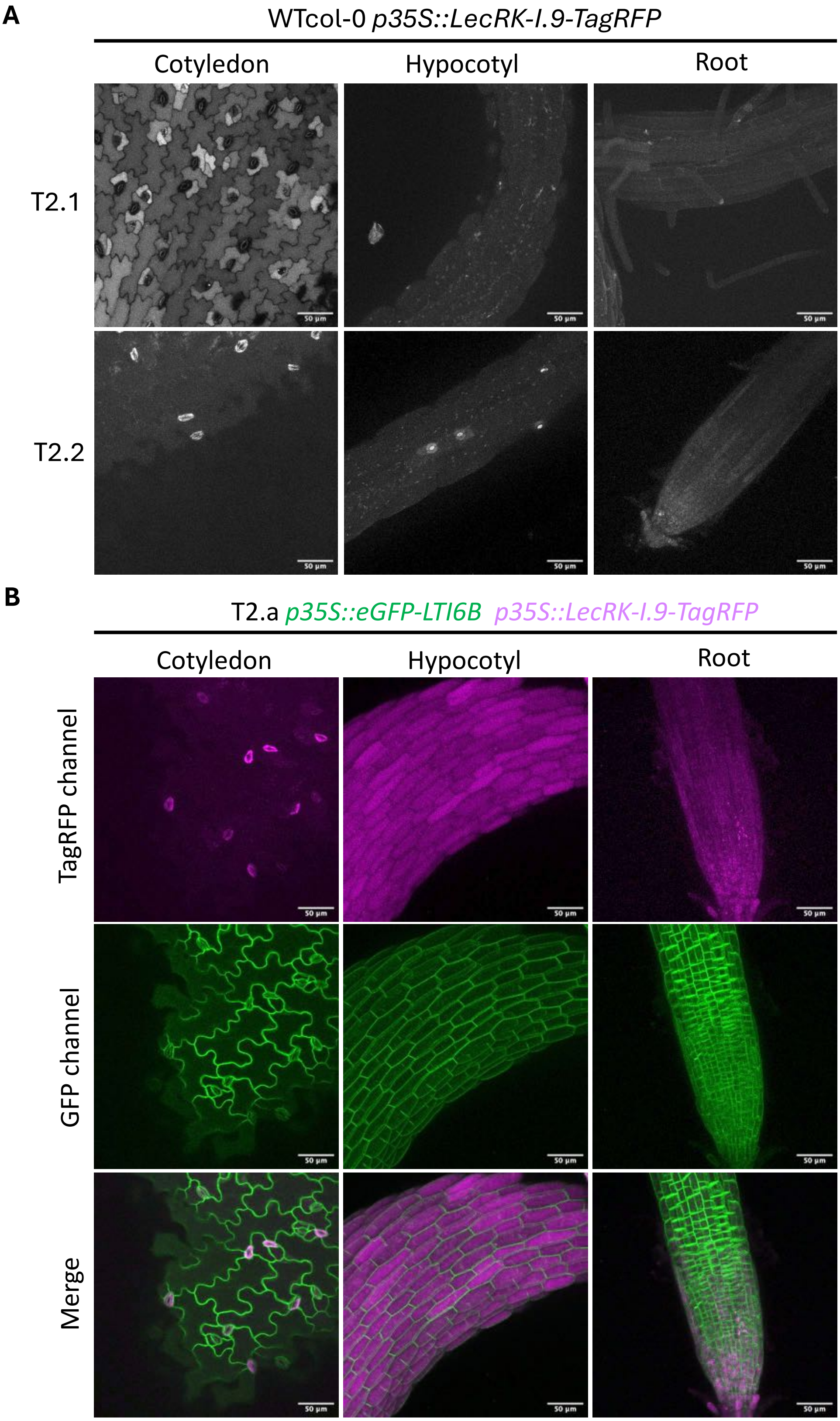


**Figure S1. Independent overexpressing lines carrying the WT version of LecRK-I.9 displayed aberrant localization of the protein and poor fluorescent signal.**

**Figure S1. Independent overexpressing lines carrying the WT version of LecRK-I.9 displayed aberrant localization of the protein and poor fluorescent signal. A.** Two independent Arabidopsis transgenic lines in WTCol-0 background (T2 generation) expressing the wild type version of LecRK-I.9 (*p35S::LecRK-I.9-TagRFP* in grey). Maximal z-stack projections showing weak or mislocalized fluorescent signal in cotyledon, hypocotyl and root of 7-days-old seedlings. Scale bar = 50 μm. (Brightness and contrast adjusted in all the images for better observation). **B.** Independent Arabidopsis transgenic line expressing simultaneously the wild type version of LecRK-I.9 (*p35S::LecRK-I.9-TagRFP* in magenta) and the membrane marker LTI6B (*p35S::eGFP-LTI6B* in green). Merge images reveal no colocalization between LecRK-I.9 and LTI6B in cotyledon, hypocotyl or root of 4-days-old seedlings. Scale bar = 50 μm. (Brightness and contrast adjusted in all the images for better observation).

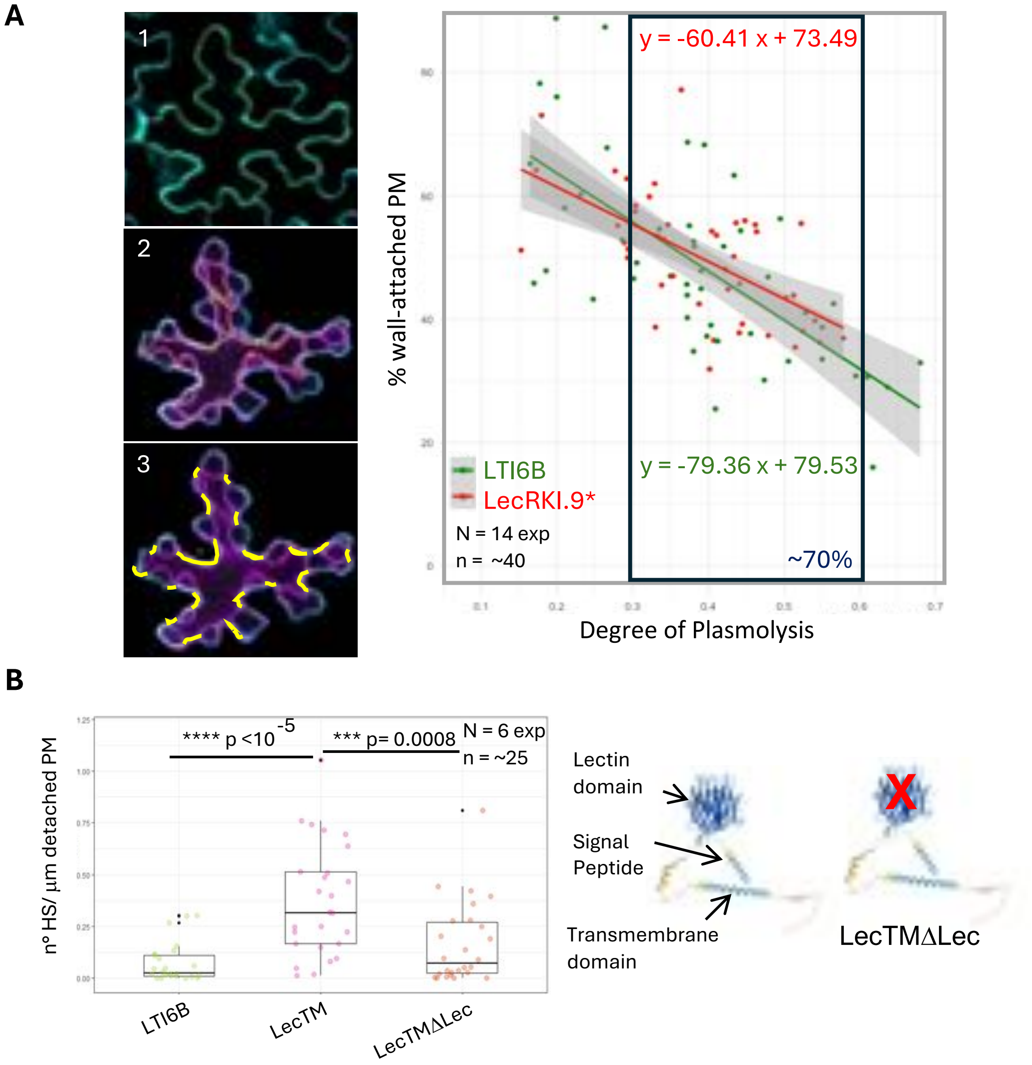


**Figure S2. LTI6B and LecRK-I.9* cells show similar plasmolysis degrees. Deletion of the lectin domain in LecTM does not enhance Hechtian strand formation. A.** Linear regression of percentage of wall-attached plasma membrane vs. degree of plasmolysis. 1. Confocal image at the middle plane of pavement cells showing the walls stained with calcofluor (cyan). Measurement of perimeter and area of the cell (yellow). 2. Segmentation of individual plasmolyzed cells (magenta) and measurement of area after plasmolysis (yellow). 3. Selection of ROIs (yellow) corresponding to segments of plasma membrane attached to the wall. The model indicates a strong negative correlation between membrane attachment and plasmolysis degree ($p < 0.001$, adjusted $R^2 = 0.413$). **B.** Alphafold-predicted structure of LecTM from Uniprot highlighting the protein domains and the deletion of the lectin. The boxplot displays the number density of Hechtian strands (HS), expressed as number of HS per µm of detached plasma membrane (PM), for LecTM, LecTMΔLec and the control LTI6B. Statistical significance was determined using the Kruskal–Wallis test followed by Dunn's post hoc test with Benjamini–Hochberg correction. No significant difference was observed between LTI6B and LecTMΔLec (*p*-value = 0.07). Sample size (N = number of experiments, n = number of images with 1 -3 cells averaged) and *p*-values are indicated on the plot.

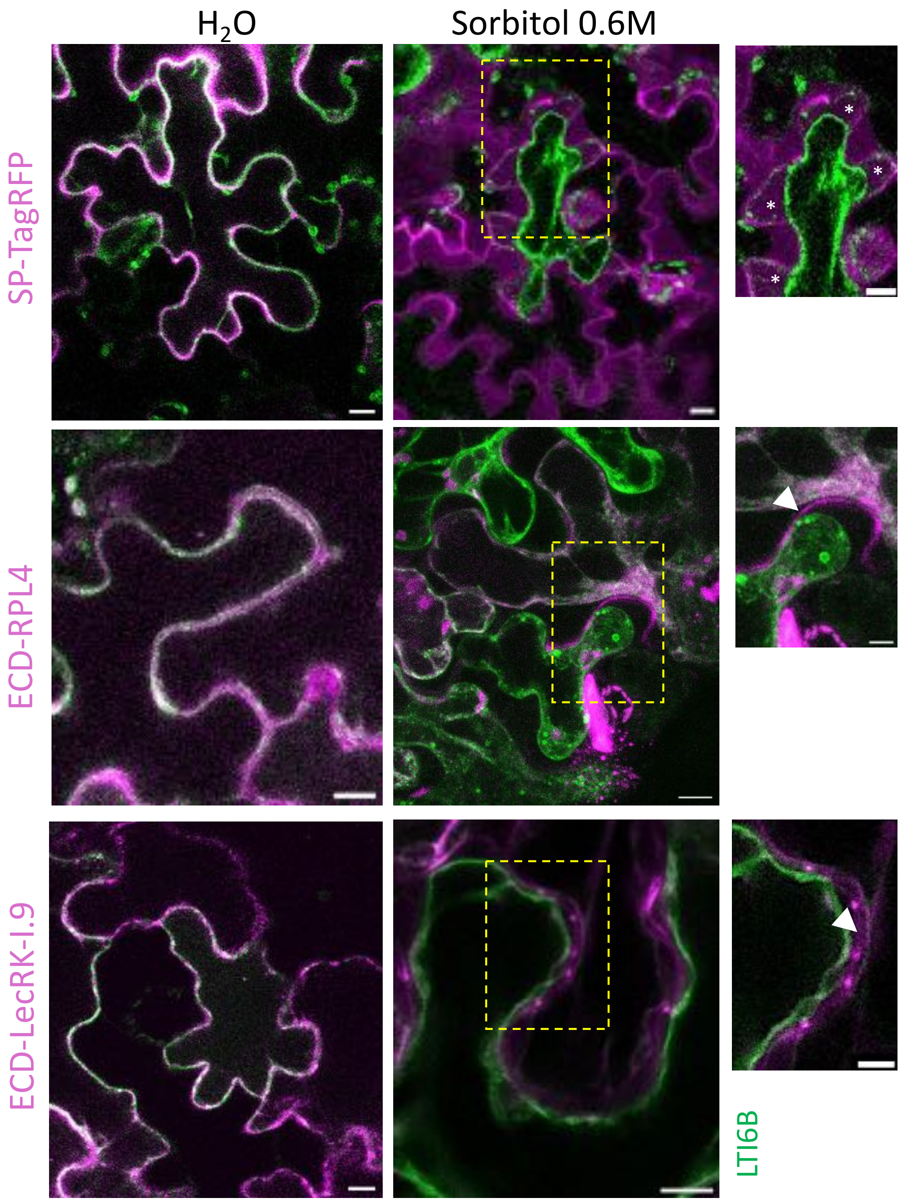


**Figure S3. The extracellular lectin domain of LecRK-I.9 (ECD-LecRK-I.9) remains in the wall during plasmolysis.** Transient transformation of *N. benthamiana* followed by plasmolysis of pavement cells with sorbitol 0.6M. In green, LTI6B-mCitrine labelling the plasma membrane (and some cytoplasmic streaming). In magenta, the negative control SP-TagRFP that cannot bind the wall; the positive control ECD-RLP4 (extracellular domain of Receptor-like protein 4) fused to TagRFP that was reported to bind the wall; and ECD-LecRK-I.9 fused to TagRFP. Merged images in water and sorbitol 0.6M. Scale bars = 10 µm. Insets corresponding to the yellow ROIs, showing SP-TagRFP diffusion into the plasmolyzed space (*); and the persistent presence of ECD-RLP4 and ECD-LecRK-I.9 in the wall (arrowheads). Scale bars = 5 µm. (Brightness and contrast adjusted in all the images for better observation).

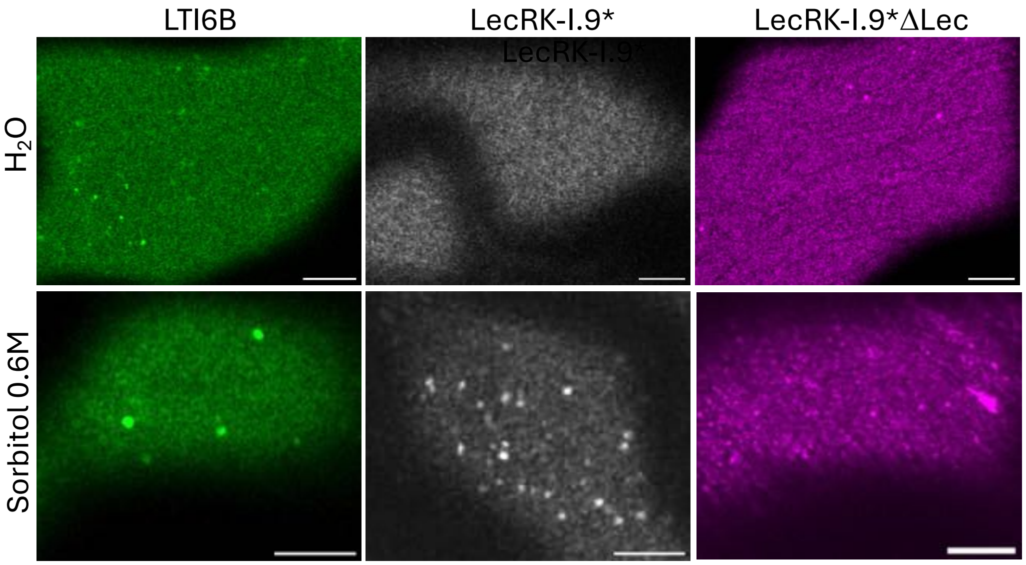


**Figure S4. Hyperosmotic stress triggered clustering of proteins. A.** Airyscan high-resolution single-plane images of cotyledon pavement cells surfaces showing the plasma membrane distribution of LTI6B (green), LecRK-I.9* (grey) and LecRK-I.9*Δlec (magenta) in water and after 30 minutes of sorbitol 0.6M incubation, condition in which clusters are observed. Scale bar = 5 µm. (Brightness and contrast adjusted in all the images for better observation).

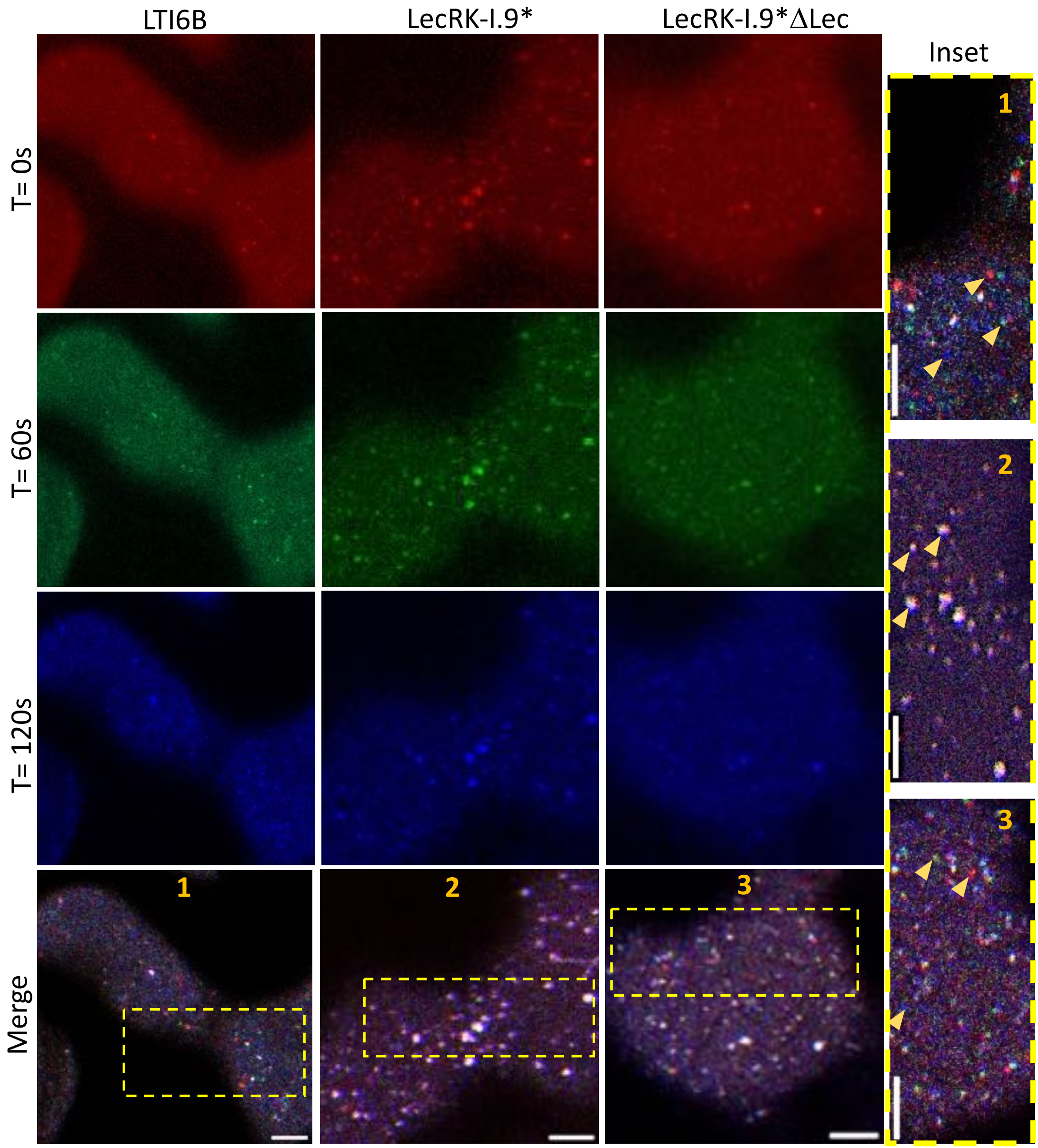


**Figure S5. LecRK-I.9* overexpressors in Arabidopsis displayed clusters under milder hyperosmotic treatments, which are less mobile than LecRK-I.9*ΔLec clusters.** 2-minutes time-lapse series of cotyledon pavement cells surfaces showing the mobility of clusters after 30-minutes of sorbitol 0.3M incubation. Frames are color-coded: Time = 0 seconds (red), Time = 60 seconds (green), and Time = 120 seconds (blue). Clusters that did not move appear white in the merge images and the ones that move appear in colors (yellow arrowheads). 1., 2., and 3. indicate the insets of LTI6B, LecRK-I.9* and LecRK-I.9*ΔLec respectively (yellow). Scale bar = 5 μm. (Brightness and contrast adjusted in all the images for better observation).

**A**

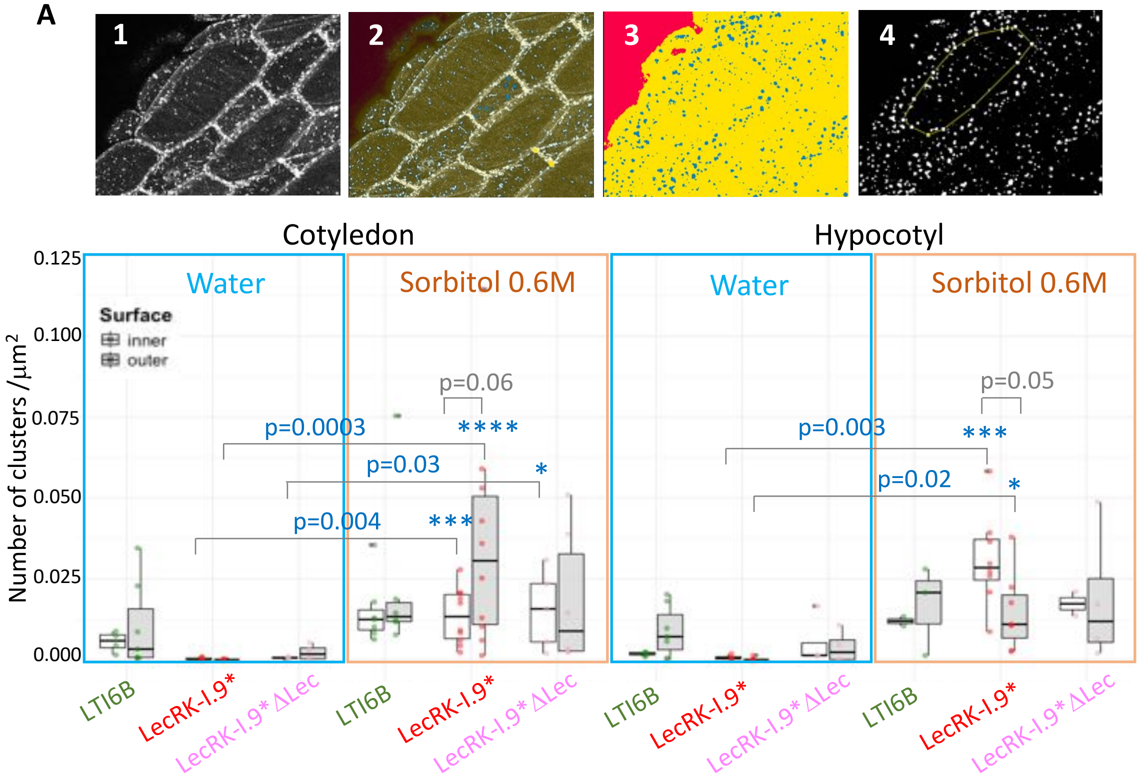


**B**

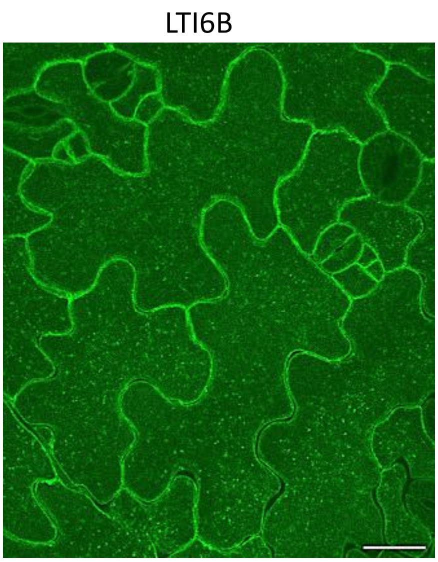


**C**

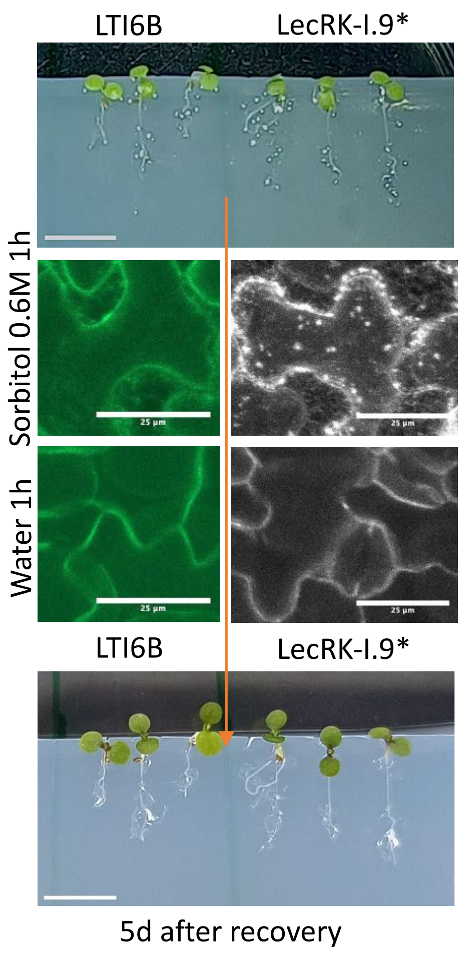


**Figure S6. Hyperosmotic treatments triggered LecRK-I.9* clustering mainly in the outer surface of cotyledons epidermal cells and inner surface of hypocotyls. The asymmetrical pattern is lost without the lectin domain.**

**Figure S6. Hyperosmotic treatments triggered LecRK-I.9* clustering mainly in the outer surface of cotyledons epidermal cells and inner surface of hypocotyls. The asymmetrical pattern is lost without the lectin domain. A.** Quantification of number of clusters per area, on the inner and outer surfaces of epidermal cells in cotyledons and hypocotyls. 1. Example of an image showing the outer surface of a hypocotyl with clusters. 2. Training of the classifier. 3. Segmentation of the image in Clusters (blue), Plasma Membrane (yellow) and Background (red). 4. Mask (black) plasma membrane and background, and selection of ROIs (yellow). The boxplots display the density of clusters, expressed as number of clusters per $\mu m^2$, for LTI6B, LecRK-I.9* and LecRK-I.9*ΔLec. Water vs. sorbitol 0.6M conditions were compared and statistical significance was determined using the Kruskal–Wallis test followed by Dunn's post hoc test with Benjamini–Hochberg correction. Inner vs. Outer surfaces within each genotype were compared by Wilcoxon test. N = 3 experiments, n = ~3-9 plants/genotype/treatment (average 1-3 cells per image). *p*-values are indicated on the plots. Asterisks denote significant differences. **B.** Confocal image showing 7 days-old Arabidopsis cotyledon pavement cells with clusters (LTI6B signal in green), that were able to maintain most of their volume despite the 1-hour incubation with sorbitol 0.6M. Scale bar = 20 μm. (Brightness and contrast adjusted for better observation). **C.** 4-days-old LTI6B and LecRK-I.9* overexpressing lines were imaged after incubation for 1 h with sorbitol 0.6M; and after recovery in water for 1h (Scale bars = 25μm). Then, the seedlings were kept in a growing chamber where they survived at least 5 days. Scale bar = 5mm.

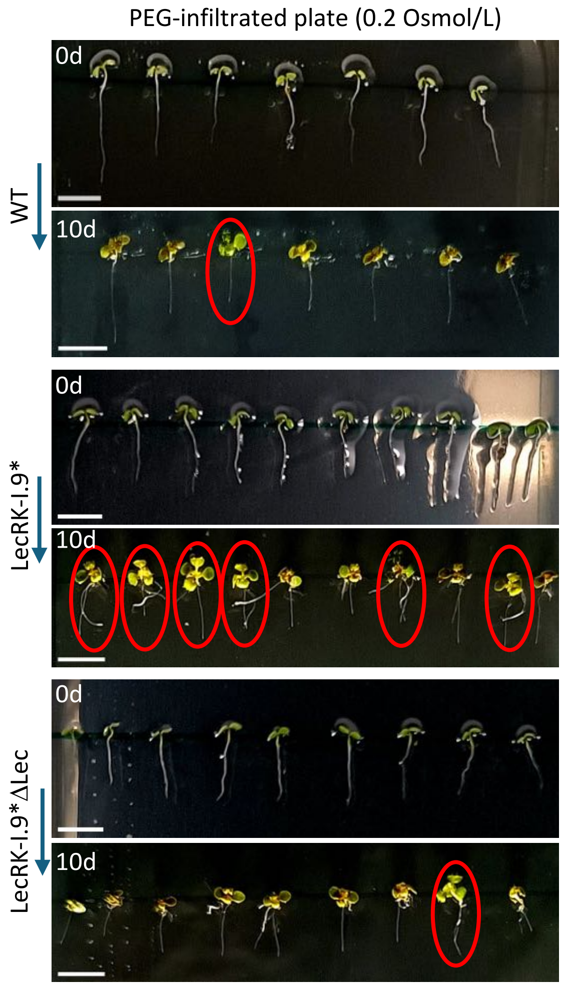


**Figure S7. LecRK-I.9* presented higher frequency of seedlings that were resistant to low water potential conditions.** Images depicting 5-d-old WT, LecRK-I.9* and LecRK-I.9*ΔLec seedlings that were immediately transferred to PEG-infiltrated plates (0d); and the same seedlings after 10 days (10 d) in those plates. The seedlings that were able to expand their cotyledons and leaves considerably were considered as resistant to low water potential conditions (circled in red). Scale bars = 5mm.

| Construct | Primer | Sequence 5' - 3' | |
|---|---|---|---|
| *p35S::LecRK-I.9-TagRFP* | Cloning | F | AAAAAGCAGGCTTCATGGCTCGTTGGTTGCTT |
| | | R | AGAAAGCTGGGTCCCTCTGACTGCTGATGCTG |
| | Sequencing | F | GCTGATTGGTTCTATAAGAAATCTAGTATTTTC |
| | | R | CGCTCACCAAACATAGAAATGC |
| | | F | TCTGTTTGCTGGGTTCTCTG |
| | | F | ATGGCCAGGTTTCATGAG |
| *p35S::ECD-LecRK-I.9-TagRFP* | Cloning | F | AAAAAGCAGGCTTCATGGCTCGTTGGTTGCTT |
| | | R | AGAAAGCTGGGTCAGATACTTTCTTATGTGGAG |
| *p35S::LecRK-I.9*-TagRFP* | Mutagenesis | F | AGCTGCATTTGGAGAAGTTTATAGAGG |
| | | R | GCTGCAGCAAATTCATCTTTGCTGAACC |
| | Sequencing | F | TCTGTTTGCTGGGTTCTCTG |
| | | F | ATGGCCAGGTTTCATGAG |
| | | F | GCTGATTGGTTCTATAAGAAATCTAGTATTTTC |
| | | R | CGCTCACCAAACATAGAAATGC |
| *p35S::LecRK-I.9* ΔLec-TagRFP* | Mutagenesis | F | CAGCGACTTGATATCTCAAAAC |
| | | F | ACTTGATACAGAGCTCAG |
| *p35S::LecTM-TagRFP* | Cloning | F | AAAAAGCAGGCTTCATGGCGGTCTTCAAAACC |
| | | R | AGAAAGCTGGGTCCTTCTTGACTTCTTCTTCAA |
| | Sequencing | F | GCTGATTGGTTCTATAAGAAATCTAGTATTTTC |
| | | R | CGCTCACCAAACATAGAAATGC |
| *p35S::LecTM ΔLec-TagRFP* | Mutagenesis | F | GATCCAAACGACGTTTCC |
| | | R | ATTTACATCTGTAACAGCC |
| *p35S::SP-TagRFP-AGP18_noSP* | Cloning | F | AAAAAGCAGGCTTCCAATCTCCTATCTCTTCTCCGACC |
| | | R | AGAAAGCTGGGTC**TTA**GAATGCCATAACGAGAACGGCC |
| | Sequencing | F | GCTGATTGGTTCTATAAGAAATCTAGTATTTTC |
| | | R | CGCTCACCAAACATAGAAATGC |
| *p35S::SP-TagRFP-COBRA_noSP* | Cloning | F | AAAAAGCAGGCTTCGATCCAGAAGGCAACATTAC |
| | | R | AGAAAGCTGGGTC**TTA**GGCAGAGAAGAAGAAAAAG |
| | Sequencing | F | GGACTCCTGATGGCTATGTT |
| | | F | CATTACGTGCACATACTCGC |
| | | F | GCTGATTGGTTCTATAAGAAATCTAGTATTTTC |
| | | R | CGCTCACCAAACATAGAAATGC |
| *p35S::SP-TagRFP-LecGPI_noSP* | Cloning | F | AAAAAGCAGGCTTCTTTCTCTACCACGGATTTTTC |
| | | R | AGAAAGCTGGGTC**TCA**TATCATAAAGATCATTATTGCA |
| | Sequencing | F | GCTGATTGGTTCTATAAGAAATCTAGTATTTTC |
| | | R | CGCTCACCAAACATAGAAATGC |
| *p35S::AT14A-TagRFP* | Cloning | F | AAAAAGCAGGCTTCATGGTGCTATCCAAAGAGAAT |
| | | R | AGAAAGCTGGGTCTTTTCCAGAACCAGTGATCT |
| | Sequencing | F | GCTGATTGGTTCTATAAGAAATCTAGTATTTTC |
| | | R | CGCTCACCAAACATAGAAATGC |
| *p35S::AFL1-TagRFP* | Cloning | F | AAAAAGCAGGCTTCATGGCGTTATCCAAAGATTTG |
| | | R | AGAAAGCTGGGTCACGGTTGATCTTTTGCAGAA |
| | Sequencing | F | GCTGATTGGTTCTATAAGAAATCTAGTATTTTC |
| | | R | CGCTCACCAAACATAGAAATGC |
| *p35S::ATHK1-TagRFP* | Cloning | F | AAAAAGCAGGCTTCATGCGAGGAGATAGCTTC |
| | | R | AGAAAGCTGGGTCAGCGGACAATGAAGTTTG |
| | Sequencing | F | GCTGATTGGTTCTATAAGAAATCTAGTATTTTC |
| | | R | CGCTCACCAAACATAGAAATGC |
| | | F | TTCCACCGGATGATCTGATC |
| | | F | CTCAAGAAGAACTTGTGGAG |
| | | F | CTACAGCTCTCCTTCGGC |
| | | F | GATCCTTCTACCACTCGG |
| *p35S::MSL10-TagRFP* | Cloning | F | AAAAAGCAGGCTTCATGGCAGAACAAAAGAGTAG |
| | | R | AGAAAGCTGGGTCGTTCTTCTTTGTGAGATTAATG |
| | Sequencing | F | TGGATATTGTTGTTCAACCACG |
| | | F | GTTAAGGTGTACACGAGCC |
| | | F | GCTGATTGGTTCTATAAGAAATCTAGTATTTTC |
| | | R | CGCTCACCAAACATAGAAATGC |
| *pEAQ-HT-DEST1 ΔP19* | Mutagenesis | F | GGCGCGTGGCCGGCCTAC |
| | | R | CGAAATTACCCTTTGTTGAAAAGTCTCAATAGCCCTTTGGTCTTCTG |
| | Sequencing | F | TGATCTGCAACTTCAAGACC |
| | | R | ATGGTTTCTGACGTATGTGC |
| | | F | GCTGATTGGTTCTATAAGAAATCTAGTATTTTC |
| | | R | CGCTCACCAAACATAGAAATGC |
| *pDONR207* | attB1 | F | GGGGACAAGTTTGTACAAAAAAGCAGGCTTC |
| | attB2 | R | GGGGACCACTTTGTACAAGAAAGCTGGGTC |

**Table S1. Constructs and primers used for cloning, sequencing, and mutagenesis.**

**Table S1. Constructs and primers used for cloning, sequencing, and mutagenesis.** The first column lists all generated constructs. The second column describes the purpose of each primer (cloning, sequencing, or mutagenesis). The third column provides the forward and reverse primer sequences (5'–3'). Nucleotides underlined indicate regions overlapping with attB sites used for BP recombination into the *pDONR207* entry vector. Bold nucleotides denote stop codons included in constructs with N-terminal TagRFP fusion.
